# WWOX deficiency uncovers a cell-autonomous mechanism impairing myelin repair

**DOI:** 10.1101/2025.11.22.689900

**Authors:** Baraa Abudiab, Carlo Manenti, Srinivasarao Repudi, Sara Abu-Swai, Rania Akkawi, Takwa Jbara, Jose Davila-Velderrain, Rami I. Aqeilan

## Abstract

Remyelination is essential for neuronal function and plasticity, and its failure contributes to multiple sclerosis (MS) and other neurodegenerative disorders. Yet, the molecular programs governing oligodendrocyte precursor cell (OPC) differentiation and remyelination remain incompletely defined. Here, we identify the WW domain–containing oxidoreductase (WWOX) as a critical cell-autonomous regulator of oligodendrocyte differentiation and myelin repair. Reanalysis of single-nucleus RNA sequencing from MS lesions revealed WWOX as one of the most significantly dysregulated oligodendroglial genes. Conditional deletion of *Wwox* in oligodendroglia impaired OPC differentiation, favouring aberrant proliferation and blocking myelin regeneration after cuprizone-induced demyelination. Single-nucleus transcriptomics confirmed profound transcriptional reprogramming in WWOX-deficient oligodendroglia during remyelination, with enrichment of WNT and TGFβ signalling and cell cycle programs. Mechanistically, WWOX physically interacts with the master transcription factor SOX10 via its WW1 domain, stabilising SOX10 protein and sustaining its downstream myelin gene network. Loss of WWOX reduced SOX10 stability and activity, providing a direct mechanistic link to defective OPC differentiation. Together, our findings uncover WWOX as an essential orchestrator of remyelination and position the WWOX–SOX10 axis as a promising therapeutic target for enhancing myelin repair in MS and related demyelinating disorders.

## Introduction

Myelin is the vertebrate solution to the challenge of achieving rapid signal conduction along narrow axons^1,2^. By electrically insulating axonal segments with myelin, oligodendroglial cells enable rapid and efficient neuronal signaling via saltatory conduction^3^. Because of this fundamental role, myelin formation and maintenance are critical for the development and preservation of sensory, motor, and cognitive functions. In the central nervous system (CNS), myelin is generated by oligodendrocytes, whose membrane wraps repeatedly around axons in a tightly regulated process known as myelination. This process is developmentally protracted in humans and adaptively implicated in learning and memory^4–8^. Diverse forms of myelin damage can lead to demyelination, and myelin sheaths degenerate with age^9^. Therefore, regenerative remyelination responses are required to ensure long-term axonal integrity. Loss of efficient remyelination is considered one of the primary mechanisms underlying progressive demyelinating disease^10,11^. The production, maintenance, and repair of myelin has been proposed to underlie the vulnerability to highly prevalent human neuropsychiatric and degenerative disorders^8,12^.

Despite recent advances in the cellular and molecular mechanisms underlying successful or impaired remyelination^10^, important questions remain. CNS remyelination can be highly efficient in animal models and in specific clinical contexts, but it is often inefficient in demyelinating diseases like multiple sclerosis (MS)^13^. It can be mediated by oligodendrocyte precursor cells (OPCs) that generate new oligodendrocytes (Oli) or possibly by pre-existing oligodendrocytes. The consensus from rodent studies is that in most cases remyelination is mediated by new oligodendrocytes^13,14^. But recent evidence from studies applying single-nucleus RNA sequencing (snRNA-seq) and C-based birth-dating to autopsy samples indicate pre-existing oligodendrocytes may play a role in remyelination in the human brain^15–17^. Determining what drives regenerative remyelination, why it fails in demyelinating disease, and how it might be enhanced therapeutically is a major challenge given the rise in disability within ageing populations^10^. Because causal mechanisms can only be investigated in experimental models, integrating human data with multiple experimental paradigms can help identify and contextualize the role of underlying cellular and molecular mechanisms.

WWOX has been previously associated with CNS development and hypomyelinating disorders, including WOREE syndrome^18^ and spinocerebellar ataxia^19^. In this study, we identified WWOX as a candidate gene implicated in promoting oligodendrocyte differentiation and remyelination with implications in both clinical and experimental contexts. We analyzed Genome-Wide Association Studies (GWAS) and snRNA-seq data to identify candidate genes influencing inadequate remyelination in MS and combined in vivo and ex vivo genetic models to investigate a functional role in oligodendroglia. We mechanistically show that WWOX promotes remyelination by stabilising SOX10, a transcriptional regulator of OPC differentiation.

## Results

### WWOX is a candidate gene implicated in MS remyelination via oligodendroglia

One possible way of identifying candidate genes and biological processes influencing inadequate remyelination in demyelinating disease is by combining evidence of genetic risk and cell-type-specific expression^20–22^ (**Fig. 1a**). Using MS as a case study, we first derived gene-level association scores from the International MS Genetics Consortium GWAS dataset^23^ (**Supplementary Table S1**), treating these scores as evidence for each gene’s contribution to MS risk. To determine the cellular context in which these genes might collectively operate, we reanalyzed existing snRNA-seq data of the white matter of post-mortem tissue of individuals with progressive MS and human controls (CTR)^15^. We annotated this data to identify all major cell types, including all expected glial (astrocytes AQP4+, OPCs VCAN+, oligodendrocytes PLP+, microglia PLXDC2+) and neuronal (excitatory neurons SYT1+SNAP25+, inhibitory neurons SYT1+GAD1+) types (**Fig. 1b**, **Supplementary Fig. 1b-d**). Using data from control samples, we tested whether risk-conferring genes tend to be preferentially expressed in particular cell types, a common proxy for the relevance of that cell type in influencing risk^24^ (Methods). We found evidence of common variant GWAS risk converging to two main CNS cell types: microglial and oligodendroglia (**Fig. 1c**). A dominant microglia association is consistent with findings of the original study^23^ and with the known association of implicated genes in immunological processes^25^. The association of oligodendroglia suggests that some of the risk might be mediated by cell-autonomous processes within this lineage. To prioritize specific genes for follow-up experimental interrogation, we combined information from previously prioritized genes based on proximity to risk variants^23^, cell-type preferential expression, gene-level GWAS risk scores, and MS differential expression. We used the full snRNA-seq data to estimate expression changes in MS lesion vs CTR in the prioritized cell types (**Fig. 1d**). Among risk genes preferentially expressed in microglia we recovered known risk factors with immune function (e.g., CD58, SP140, MERTK), demonstrating the reliability of our analysis. Among the ranked genes, JAZF1 also showed differential expression in the microglia of MS lesions. In addition to MS, this transcriptional repressor has been associated with systemic sclerosis^26^ and may contribute to multiple inflammatory diseases via its immunosuppression function. In oligodendroglia, we recovered several risk genes related to oligodendrocyte differentiation and myelination (e.g., ZNF365, BCAS1, SOX8, PPAR2). Among these genes, WWOX showed the strongest differential expression in MS lesions vs CTR (**Fig. 1d**). The snRNA-seq data includes nuclei isolated from different neuropathology-defined areas of the same MS tissue, including normal-appearing white matter (NA), active (A), chronic active (CA), chronic inactive (CI) and remyelinated (RM) lesions^15^ (**Fig. 1b**). Cell type fractions qualitatively show differences in cellular composition, with changes in fractions of immune cells and oligodendroglia in active (CA, A), inactive (CI), and remyelinating (RM) lesions (**Fig. 1e**). Examination of oligodendroglial gene expression revealed that WWOX expression was highest in RM lesions, intermediate in controls, and lowest in NA regions—suggesting that WWOX may play a role in promoting remyelination in MS (**Fig. 1f**).

**Figure 1.**
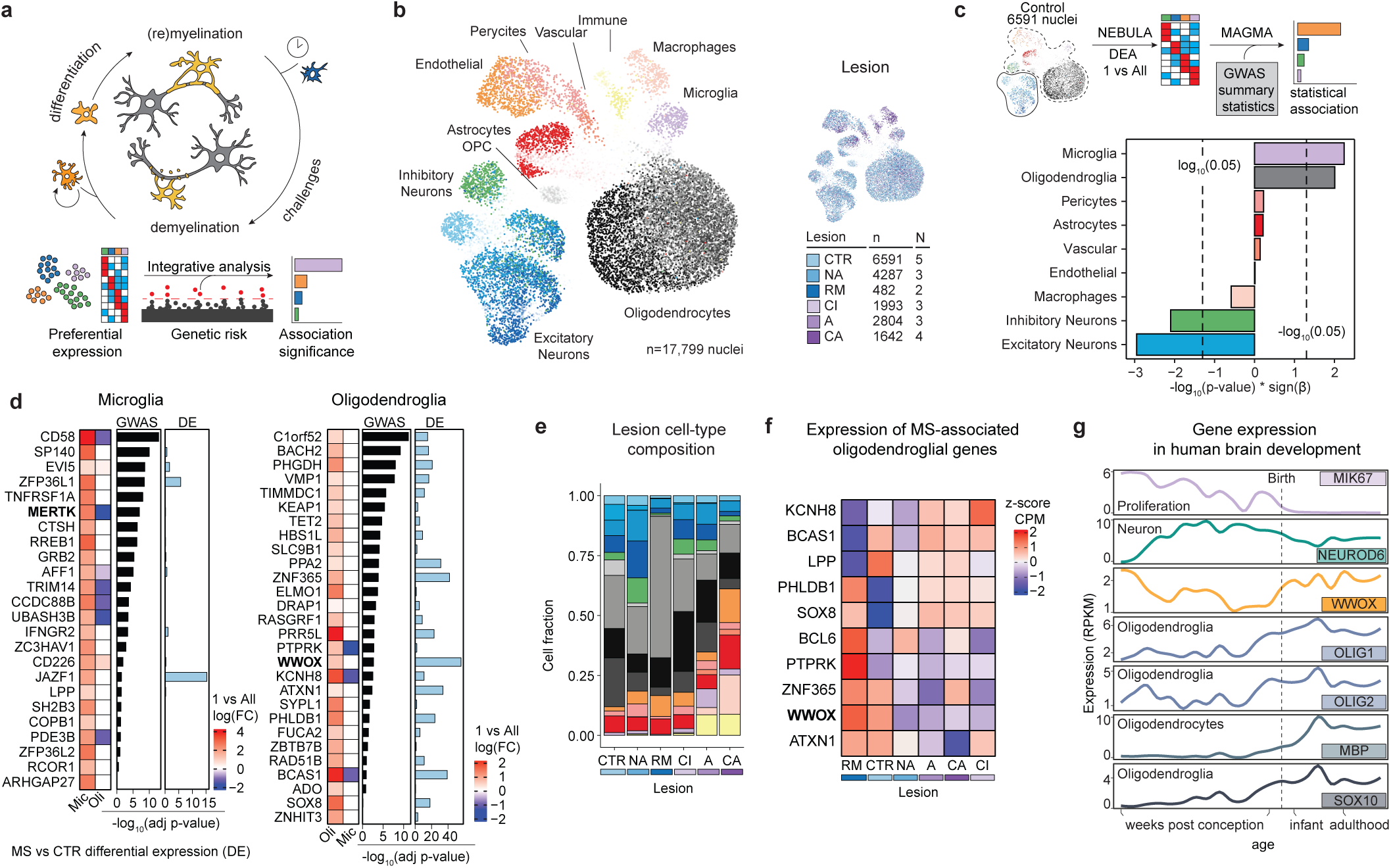
*WWOX* is implicated in MS remyelination in oligodendroglia. **a** Myelination and remyelination are essential processes for neuronal survival, function, and homeostasis of the brain. We assessed the cellular and molecular basis of impaired remyelination by relating cell-type-preferential expression and genetic predisposition to demyelinating diseases. **b** 2D representation of 17,799 nuclei from Multiple Sclerosis (MS) and control (CTR) postmortem white matter samples. Single nuclei are represented as points and labeled by cell type (left) or lesion type (right) (CTR: Control; NA: Normal Appearing white matter; RM: Remyelinating; CI: Chronically Inactive; A: Active; CA: Chronically Active). **c** Cell-type MS genetic variability association analysis. Bar lengths define the strength of the association as quantified via MAGMA (-log10(adjusted p-value)) using cell-type-preferential expression in CTR samples (NEBULA, Differential Expression Analysis (DEA) between cell types in CTR samples, adjusted p-value < 0.01). Positive (negative) bars display positive (negative) associations between cell-type-preferential expression and MS genetic variability. Vertical dashed lines highlight significance thresholds for the association (p-value < 0.05, MAGMA gene-covar, linear regression). **d** Candidate genes driving cell-type specificity and GWAS associations in microglia or oligodendroglia. Candidate genes were identified by prioritization in the original MS GWAS study and for their cell-type preferential expression. Candidate genes are ranked based on MS GWAS genetic variability (MAGMA gene-level score), and MS transcriptional variability was considered for further prioritization (NEBULA, differential expression MS vs CTR, adjusted p-value < 0.01). **e** Cell-type composition across MS lesions and CTR samples. **f** Relative expression of oligodendroglial candidate genes differentially expressed across lesions (Wilcoxon signed-rank test, log2(FoldChange) > 0.05, detected in at least 25% of the nuclei, and adjusted p-value < 0.01). **h** Expression of *WWOX* and canonical marker genes across human brain development.

The cell and context specificity of WWOX expression is also supported by reference developmental transcriptomic data of the human brain and cell-sorted mouse brain^27,28^ (**Fig. 1g**). WWOX expression follows a developmental trajectory that mirrors that of myelin marker proteins (MBP) and oligodendroglial TFs (OLIG1/2, SOX10). These genes are expressed at times overlapping oligodendrogenesis and myelination^29^. Within the oligodendroglial lineage, WWOX is expressed the highest in OPCs followed by newly formed and mature oligodendrocytes (**Supplementary Fig. 1f**). WWOX has been implicated in CNS development and severe hypomyelinating disorders, including WOREE syndrome^18^ and spinocerebellar ataxia^19^, yet its direct role in oligodendroglial biology has not been explored. Guided by these converging lines of evidence, we focused our experimental investigations on defining the cell-autonomous function of WWOX in oligodendroglia.

### WWOX deficiency impairs OPC differentiation and OL maturation in vitro

To investigate the effect of WWOX loss in OPC differentiation and maturation *in vitro*, we established enriched primary OPC cultures from Wild-Type (WT) and *Wwox*-null (KO) pups; the latter from a KO model previously generated by our group^18^ **(Fig. 2a)**. We phenotyped the cultures using RNA-sequencing at 2 days of differentiation and immunostaining at 5 and 8 days. At the early differentiation stage, KO samples expressed higher levels of Radial Glia (RG) markers, and lower levels of oligodendroglial markers relative to WT cells **(Fig. 2b)**, suggesting potentially delayed or impaired differentiation in *Wwox*-depleted oligodendroglia. Genome-wide differential expression and pathway analysis identified global changes consistent with this observation. KO cell showed reduced expression of myelin-associated genes (e.g., *Mag*, *Cnp*), together with increased expression of cell-cycle (*Cenpf, Cdk1*), proliferation (*S100a6*, *Anxa1*), and WNT-associated genes (e.g., *Wnt2*, *Fzd6*, *Ccn4*, *Ccnd1*, *Nfatc4*, *Gpc4*, *Vangl1*, *Rspo3*, Fzd7) **(Fig. 2c)**. Stem cell and cell cycle associated pathways were enriched in KO cells, while oligodendroglial signatures (Oli and OPC) and metabolic (oxidative phosphorylation, electron transport) and neuronal support pathways (glutamatergic synapse) enriched in WT cells **(Fig. 2d)**. These molecular changes were further validated at protein and cellular levels at day 5 and 8. A significantly lower fraction of cells was immunoreactive for MBP in KO cultures, while a larger fraction was immunoreactive for PDGFRα **(Fig. 2e)**. Quantification of MBP-positive cell numbers confirmed this observation, with significantly fewer MBP-positive cells in the KO group **(Supplementary Fig. 2g)**. Together with this impaired differentiation phenotype, MBP+ cells also showed morphological alterations in KO cultures. KO MBP+ cells exhibited reduced surface area and branching complexity, as quantified by Sholl analysis, suggesting potential morphological immaturity in these cells **(Fig. 2f)**. Taken together, these data imply that oligodendroglial WWOX expression is necessary for normal OPC differentiation and oligodendrocyte maturation in murine cell cultures and suggest an unforeseen cell-autonomous role.

**Figure 2.**
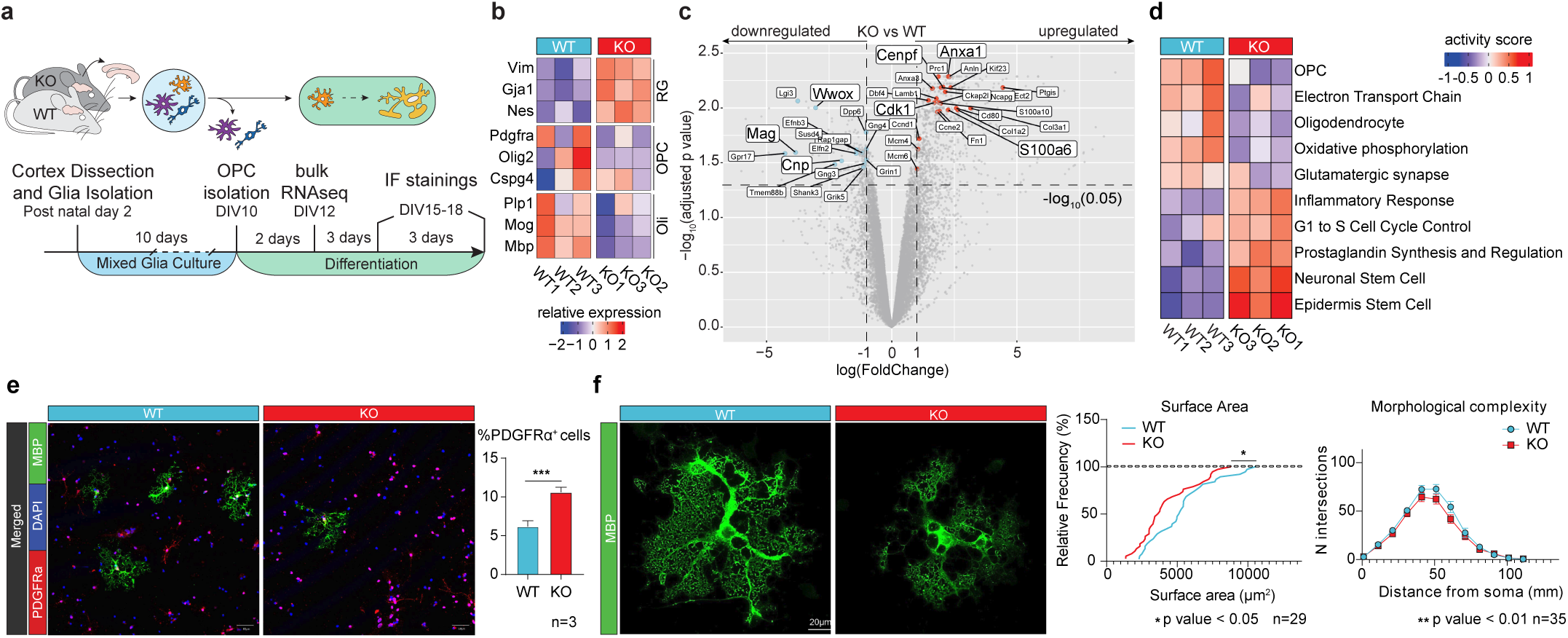
WWOX deficiency impairs oligodendrocyte differentiation and maturation. **a.** Experimental design used to assess primary cultures of Oligodendrocyte Precursor Cells (OPCs) and their differentiation (DIV: Days In Vitro). **b** Relative expression (z-score) of cell-type marker genes for Radial Glia (RG), OPCs, and Oligodendrocytes (Oli) across samples. **c** Volcano plot showing differential expression of leading edges from gene programs in **d**. For each gene program, the first five leading edges were visualized on the volcano plot if differentially expressed (adjusted p-value < 0.05 and log2(FC) > 1). Upregulated (downregulated) leading edges with respect to WWOX-deficient cultures were annotated in red (blue). All other genes, except *Wwox*, have been colored in grey even if differentially expressed. Dashed lines define differential expression thresholds for adjusted p-value and log2(Fold Change). **d** Inferred activity of selected gene programs. Gene programs were selected from the top 5 most over- and underrepresented gene programs (based on adjusted p-value) for each tested gene set (KEGG, WikiPathways, and Tabula Muris). **e** Immunofluorescence staining (IF) of OPCs (PDGFRa+) and Olis (MBP+) in WT and KO cultures (data are shown as means +/− SEM; two-tailed t-test). **f** Morphological analysis of Oli. From left to right: IF of CTR and KO Oli (MBP+); quantification of Oli surface area (Kolmogorov-Smirnov test); Sholl analysis of branching complexity based on the number of intersections with concentric circles drawn from the cell soma (data are shown as means +/− SEM, ANOVA).

### WWOX deficiency causes subtle myelination defects in vivo and increases susceptibility to age-related demyelination

To assess the role of WWOX in oligodendroglial development in vivo, we used an Olig2-Cre–driven conditional Wwox knockout (O-KO) model targeting the oligodendrocyte lineage^18^ (**Supplementary Fig. 3a**). Efficient recombination was confirmed by the marked reduction of WWOX in CC1^+^ oligodendrocytes (**Supplementary Fig. 3b**). Under baseline conditions, O-KO mice showed no differences in the proportions of PDGFRα^+^ OPCs or CC1^+^ oligodendrocytes in the corpus callosum compared with controls, indicating similar lineage cell numbers (**Supplementary Fig. 3c**). Likewise, MBP immunofluorescence across the cortex and corpus callosum was comparable between genotypes, with only minor differences detected in the cerebellum (**Supplementary Fig. 3d**), suggesting largely intact myelination.

However, transmission electron microscopy (TEM) imaging in the corpus callosum revealed a subtle reduction in myelin thickness in O-KO mice with no apparent changes in the number of myelinated axons (O-CTR: 117.6 ± 12.5 vs. O-KO: 125.1 ± 9 per FOV; P = 0.22), potentially reflecting thinly remyelinated or partially demyelinated fibers (**Supplementary Fig. 3e**). Given that remyelination efficiency naturally declines with age^10,30,31^, we hypothesized that this mild defect might predispose O-KO mice to age-related demyelination. Consistent with this idea, 18-month-old O-KO mice exhibited fewer myelinated axons than controls (O-CTR: 130.1 ± 28.6 vs. O-KO: 102.3 ± 31.8 per FOV; P = 0.0044), despite similar myelin thickness (**Supplementary Fig. 3f**), suggesting impaired remyelination capacity during aging in the absence of WWOX.

### WWOX deficiency markedly impairs remyelination after cuprizone-induced demyelination

Because the mild myelin abnormalities in WWOX-deficient oligodendroglia appeared to worsen under the physiological stress of aging—when myelin integrity naturally declines^30^, we hypothesized that an acute demyelinating challenge might unmask a more pronounced phenotype and reveal the underlying cellular defect. We therefore subjected O-KO and O-CTR mice to the cuprizone model of demyelination^32,33^, administering cuprizone for five weeks followed by two weeks of recovery (**Fig. 3a**). Both genotypes exhibited comparable demyelination, confirmed by reductions in MBP immunoreactivity and Luxol fast blue staining (**Supplementary Fig. 4a–b**). Remarkably, differences emerged during the recovery phase. O-CTR mice showed robust remyelination across cortex, cerebellum, and corpus callosum, whereas O-KO mice exhibited markedly impaired myelin restoration (**Fig. 3b**; **Supplementary Fig. 4b**). Quantification revealed significantly reduced MBP signal in O-KO mice (O-CTR = 60; O-KO = 40) (**Fig. 3c**). TEM analysis further supported impaired remyelination as revealed by thinner myelin sheaths and fewer myelinated axons in O-KO tissue (O-CTR: 59.1/FOV; O-KO: 36.1/FOV) (**Fig. 3d**).

**Figure 3.**
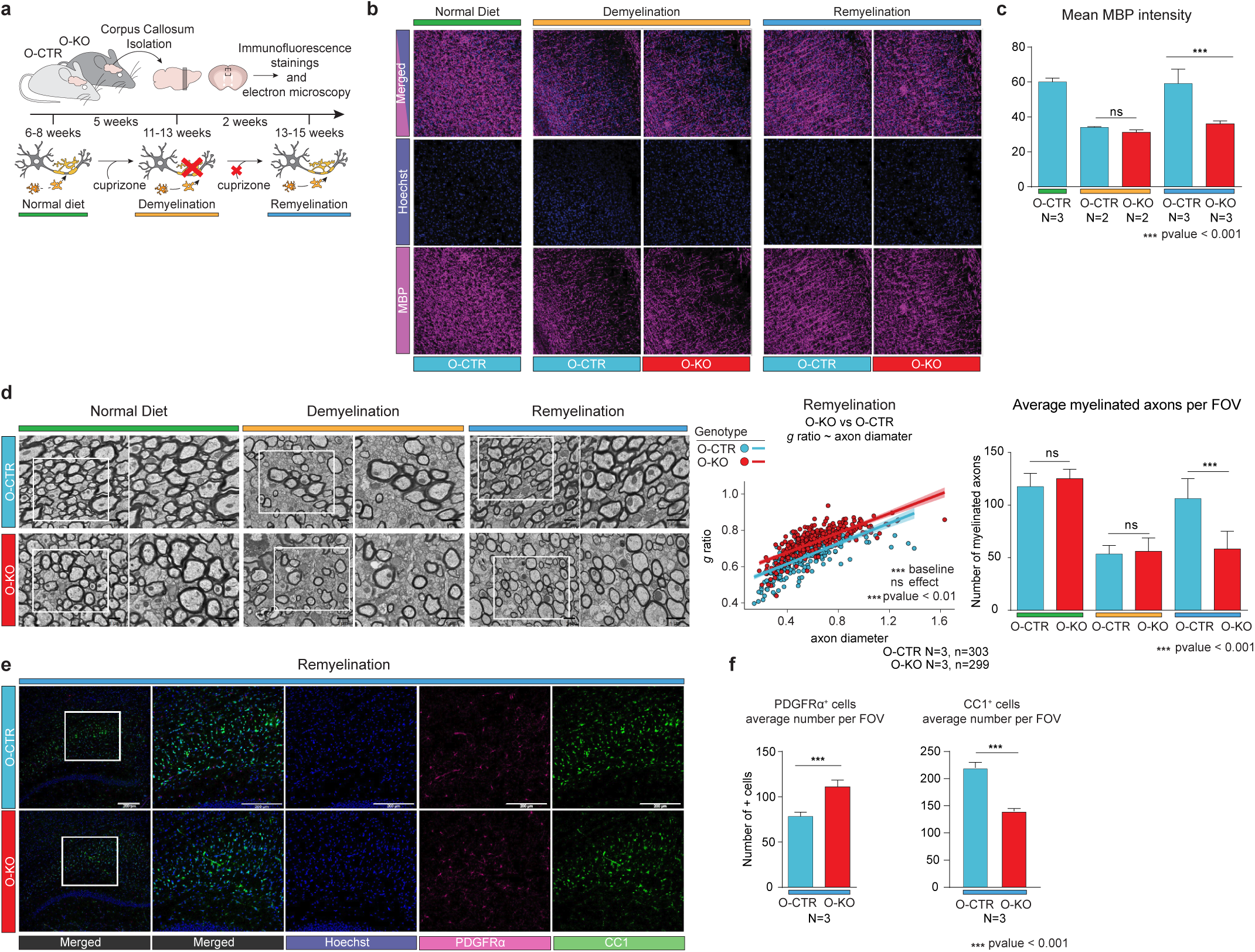
Cuprizone-induced demyelination reveals remyelination defects in *WWOX*-deficient oligodendroglia. **a** Experimental design of cuprizone-induced remyelination in mouse models. **b** Representative immunofluorescence (IF) for MBP in brain sections following demyelination and remyelination periods. **c** Quantification of MBP intensity in conditions shown in b for three anatomically matched sagittal sections per animal (data are shown as means +/− SEM; two-tailed t-test). **d** Transmission Electron Microscopy (TEM) analysis of myelin. Left: representative images of the midsagittal corpus callosum following remyelination; white squares indicate magnified regions. Middle: g-ratio analysis quantifying differences due to oligodendroglial WWOX deficiency (O-KO) in the average amount of myelin and the effect of axonal diameter (linear regression). Right: quantification of the number of myelinated axons per Field Of View (FOV) (data are shown as means +/-SEM; two-tailed t-test; n.s., not significant). e Representative IF for OPCs (PDGFRa+) and mature oligodendrocytes (CC1+) in the corpus callosum. f Quantification of PDGFRa+ and CC1+ cells in anatomically matched sagittal sections (data are shown as means +/− SEM; two-tailed t-test).

To determine whether defective OPC differentiation contributed to this remyelination failure, we quantified PDGFRα^+^ OPCs and CC1^+^ oligodendrocytes. O-KO mice showed a pronounced reduction in mature CC1^+^ cells (O-CTR: 220/FOV; O-KO: 140/FOV), accompanied by an accumulation of OPCs (O-CTR: 78.25/FOV; O-KO: 111/FOV) (**Fig. 3e–f**), consistent with a blockade in lineage progression.

Together, these findings identify WWOX as a critical cell-autonomous regulator of myelin regeneration, enabling OPCs to generate new myelinating oligodendrocytes following demyelinating injury.

### Single-cell transcriptomic analysis reveals WWOX-mediated alterations in remyelinating OPCs and OLs

To investigate the molecular alterations accompanying WWOX deficiency and their interaction with cuprizone-induced demyelination at single-cell resolution, we performed snRNA-seq of O-CTR and O-KO mouse cortices sampled during normal-diet, demyelination, or remyelination (**Fig. 4a**). We recovered 40,906 high-quality nuclei and identified seven major cell types—inhibitory and excitatory neurons, astrocytes, OPCs, oligodendrocytes (Oli), microglia, and endothelial cells—based on canonical marker genes and transcriptional similarity to a matched snRNA-seq reference^34^ (**Supplementary Fig. 5 a-e**). Among these populations, Oli exhibited the most pronounced transcriptional reorganization after cuprizone treatment, forming clearly segregated clusters in low-dimensional space, consistent with the known oligodendrocyte-selective toxicity of cuprizone. Other cell types showed minimal diet-associated shifts, with the exception of a small subset of inhibitory neurons (**Fig. 4c**). We quantified transcriptional shifts in O-KO versus O-CTR samples in each diet condition with differential abundance tests using MiloR ^35^(**Fig. 4d**).

**Figure 4.**
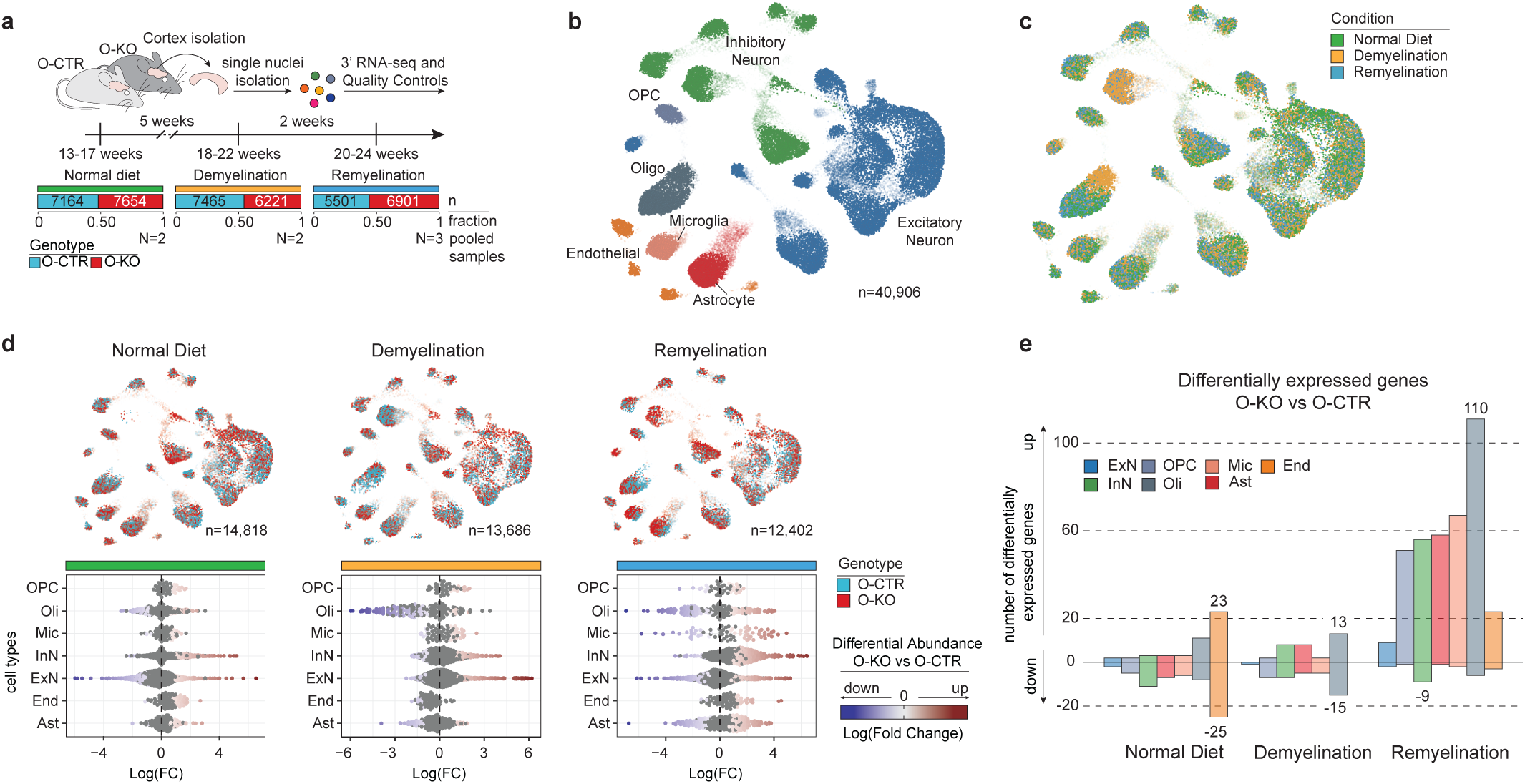
WWOX deficient oligodendroglia display transcriptional changes in remyelination. **a** Experimental design of single-nucleus RNA-sequencing (snRNA-seq) data generation in cuprizone-induced remyelination mouse models. The number of samples, and absolute number and fraction of nuclei retrieved for each combination of genotype and condition are reported. **b** 2D representation of the whole snRNA-seq dataset (40,906 nuclei) labeled by broad cell type populations. **c** 2D representation labeled by condition. **d** 2D representation with associated quantification of transcriptional shifts due to oligodendroglial *Wwox* deficiency in each condition. 2D representations are labeled by genotype. **e** Number of differentially expressed genes per cell type in each condition. (ExN: Excitatory Neurons; InN: Inhibitory Neurons; OPC: Oligodendrocyte Precursor Cells; Oli: Oligodendrocytes; Ast: Astrocytes; Mic: Microglia; End: Endothelial cells)

Under baseline or demyelinating conditions, O-KO and O-CTR samples showed only modest transcriptional differences in most cell types, apart from demyelinated oligodendrocytes and a subset of neurons. In contrast, the remyelination phase revealed widespread transcriptional divergence in O-KO cells, with oligodendroglial populations showing the strongest shifts. These global patterns matched gene-level expression changes (**Fig. 4e**): few differentially expressed genes were detected between genotypes at baseline or during demyelination, whereas remyelination was marked by a substantial upregulation of genes in O-KO samples. Across all conditions, oligodendroglial cells were the most transcriptionally impacted cell type during both demyelination and remyelination.

### WWOX loss disrupts oligodendroglial transcriptional programs and differentiation trajectories during remyelination

To dissect the molecular consequences of WWOX depletion within the oligodendroglial lineage, we examined transcriptomic dynamics across OPCs and oligodendrocytes (**Fig. 5a**). We isolated and reanalyzed 4,799 single-nucleus transcriptomes from O-CTR and O-KO mice spanning baseline, demyelination, and remyelination. The resulting landscape recapitulated the expected continuum of oligodendroglial differentiation, from OPCs to mature oligodendrocytes, and revealed a cuprizone-induced “damaged” oligodendrocyte state (**Fig. 5b**). To assess how WWOX loss affects lineage progression, we compared differentiation trajectories between genotypes using differential trajectory analysis (condiments^36^) across all conditions (**Fig. 5c**). Trajectories diverged only during the remyelination phase, with clear genotype-specific shifts in both OPCs and oligodendrocytes, while cells under normal diet or demyelination showed minimal differences.

**Figure 5.**
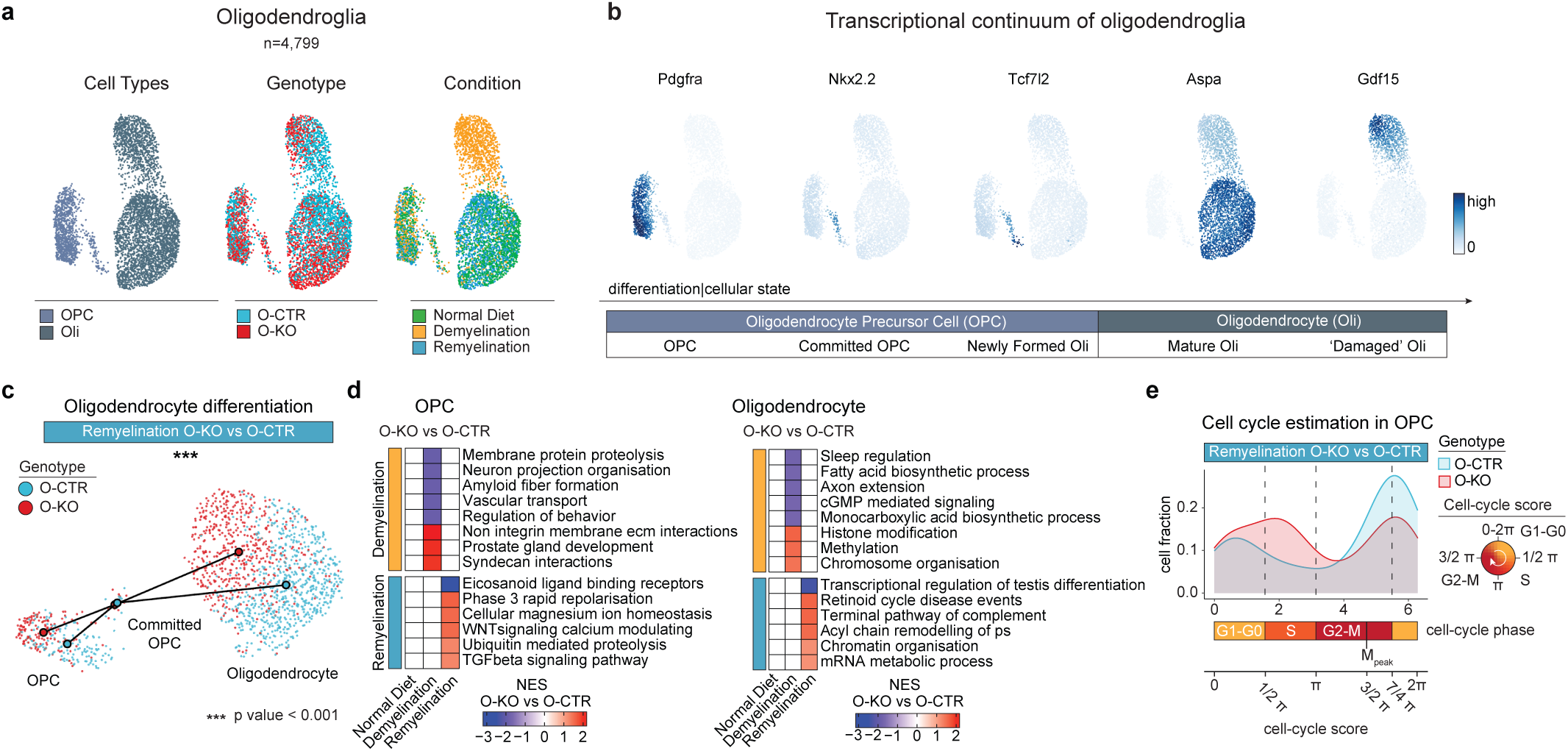
In remyelination, WWOX deficient OPCs proliferate instead of differentiate. **a** 2D representation of 4,799 oligodendroglial nuclei labelled by cell type, genotype, and condition. **b** Oligodendroglial transcriptional continuum displayed by the expression (logcounts) of selected marker genes. **c** Differential trajectory analysis. 2D representation of oligodendroglia following remyelination, overlaid by genotype-specific differentiation trajectories. **d** Inferred activity of gene programs across conditions. The 5 most over- and underrepresented gene programs are reported in each condition (ranked based on adjusted p-value). Fewer gene programs were reported if not statistically significant (adjusted p-value < 0.05) and the score displayed is the Normalized Enrichment Score (NES). **e** Fraction of OPCs for each cell-cycle phase during remyelination. Cell-cycle positions are reported as continuous (as estimated via tricycle) and discrete cell-cycle phases.

Consistent with this, differential gene expression and pathway enrichment analyses revealed no major alterations under baseline conditions (**Fig. 5d**). In contrast, during remyelination, O-KO OPCs upregulated genes associated with TGFβ and WNT/Ca²^+^ signaling—pathways known to impair myelin repair when dysregulated^37^. Oligodendrocytes from O-KO mice displayed increased expression of genes linked to chromatin organization and remodeling during demyelination and remyelination, accompanied by downregulation of metabolic and homeostatic processes such as fatty acid biosynthesis, monocarboxylic acid metabolism, and axon–glia interaction pathways (**Fig. 5d**).

Given the established roles of TGFβ and WNT signaling in controlling OPC proliferation and differentiation, we next examined cell-cycle status. Single-cell phase inference using tricycle^38^ revealed a higher fraction of O-KO OPCs in S-phase, indicating an expanded cycling population during remyelination (**Fig. 5e**). By contrast, most O-CTR OPCs resided in G1–G0, consistent with a shift toward differentiation rather than proliferation.

Together, these data demonstrate that WWOX is required to maintain proper lineage progression during myelin repair, restraining aberrant OPC proliferation while enabling the transcriptional programs necessary for successful differentiation and remyelination.

### WWOX directly binds and stabilizes SOX10, a key regulator of oligodendrocyte terminal differentiation

Given that WWOX is a scaffold adaptor protein known to interact with and modulate transcription factors (TF)^39,40^, we reasoned that its cell-autonomous and context-specific effects in oligodendroglia might arise from direct regulation of a key determinant of OPC differentiation. To identify potential candidates, we examined TF expression in WWOX-deficient versus wild-type OPCs using bulk RNA-seq from our in vitro model (**Fig 2**). Among all oligodendroglial TFs assessed, *Sox10* emerged as the only factor significantly reduced in KO cells (**Fig. 6a**). A similar reduction was detected in vivo in Wwox-deficient OPCs during remyelination (**Fig. 6b**). Immunostaining confirmed these transcriptional trends: O-KO mice displayed a substantial decrease in SOX10^+^ nuclei within the corpus callosum during remyelination (O-CTR: 1550 ± 200; O-KO: 937.8 ± 168) (**Fig. 6c**; **Supplementary Fig. 6a**). Consistent with a functional SOX10 deficit, gene-set enrichment analysis (GESA) revealed under-representation of SOX10 target genes (positive SOX10 regulon^41^) in O-KO OPCs during remyelination and in O-KO oligodendrocytes during demyelination (**Fig. 6d**).

**Figure 6.**
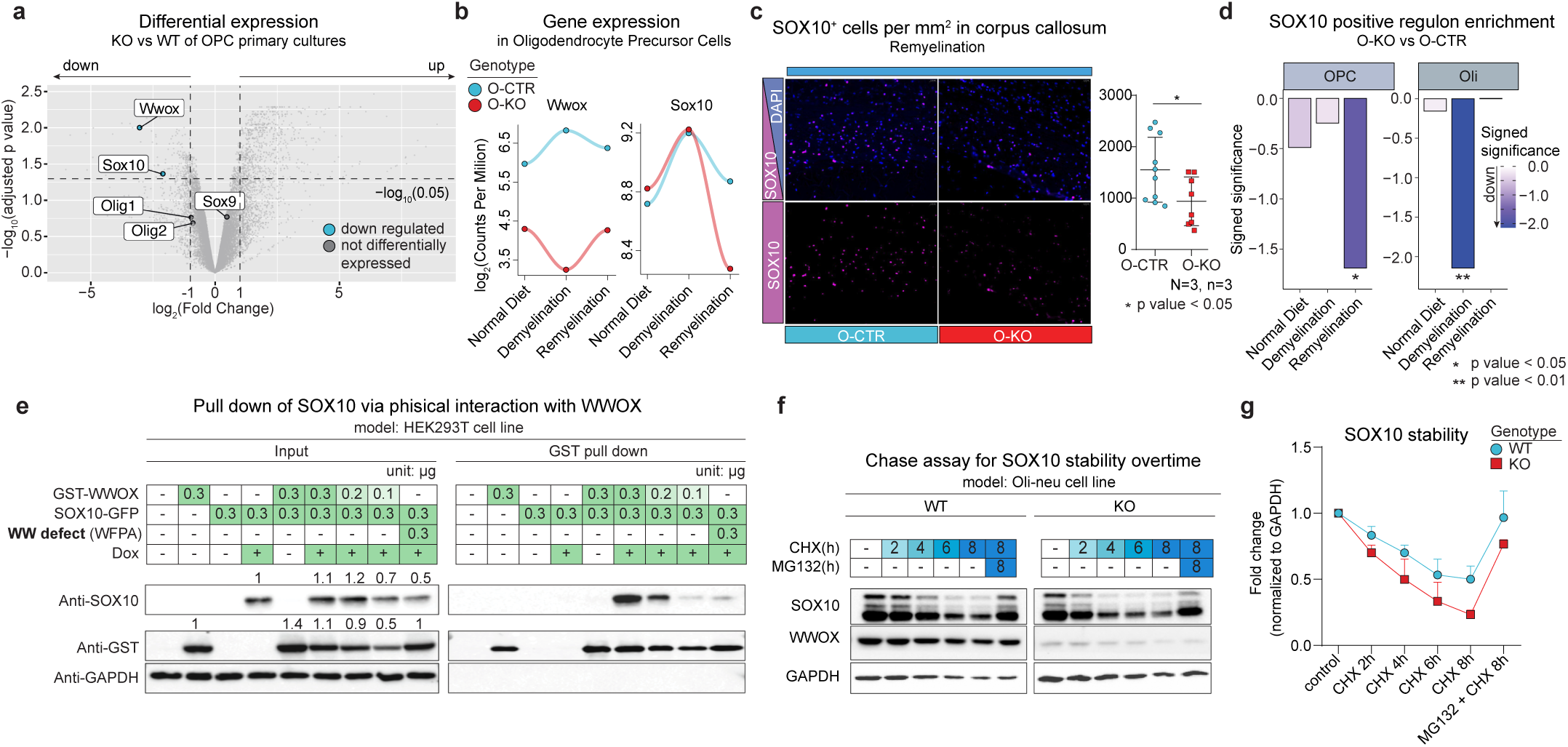
WWOX modulates SOX10 activity through direct protein-protein interaction. **a** Volcano plot showing differentially expressed oligodendroglial TFs in primary OPC cultures. **b** Expression trends of Wwox and Sox10 between KO and CTR OPCs across conditions in snRNA-seq data. **c** Representative IF and quantification of SOX10+ cells in the corpus callosum of KO and CTR mice following remyelination (data are presented as mean ± SEM; two-tailed t-test). **d** Gene set enrichment analysis of the SOX10 positive regulon in OPCs and oligodendrocytes across conditions in snRNA-seq data. **e** Western blots of GST pull-down assay for physical interaction of WWOX and SOX10 in the HEK293T cell line. The role of the WW domain in the physical interaction was tested with the WFPA mutant WWOX protein carrying a defect in the WW domain. **f** Western blots of Cycloheximide (CHX) chase assay of SOX10. Cells were subjected to a protein synthesis inhibitor, CHX (20 μg/ml), to measure SOX10 stability in both *Wwox* KO and CTR immortalised OPC cell lines (Oli-neu). Proteasome inhibitor MG132 (10 μM) was used to assess proteasome-mediated degradation of SOX10. **g** Quantification of SOX10 protein levels from CHX chase assay for each genotype.

SOX10 contains a PPxY motif, a canonical binding site for the WW1 domain of WWOX (**Supplementary Fig. 6b**), suggesting a potential physical interaction. Co-immunoprecipitation experiments using cortical lysates from a newly generated HA-tagged WWOX knock-in mouse confirmed an endogenous WWOX–SOX10 interaction (**Supplementary Fig. 6c**). A GST pull-down assay further validated this binding and demonstrated dose-dependent enhancement of SOX10 levels with increasing WWOX expression (**Fig. 6e**). Crucially, a WW1-domain mutant of WWOX (WWOX-WFPA^42^) failed to bind SOX10 and did not result in increased SOX10 protein level, indicating that the WW1 domain is essential for this interaction (**Fig. 6e**; **Supplementary Fig. 6b**).

To determine whether WWOX affects SOX10 stability, we measured SOX10 half-life in a CRISPR-engineered WWOX KO OPC line (Oli-neu). Following cycloheximide (CHX) treatment, WWOX-deficient cells displayed accelerated SOX10 decay compared to WT controls, demonstrating that WWOX prolongs SOX10 stability (**Fig. 6f–g**). Proteasome inhibition with MG132 rescued SOX10 levels in both genotypes, confirming that SOX10 undergoes proteasome-dependent turnover and supporting a stabilizing role for WWOX. Given the enrichment of ubiquitination-related pathways in our snRNA-seq data, we further examined SOX10 ubiquitination. Immunoprecipitation revealed a modest increase in ubiquitinated SOX10 in WWOX-deficient cells (**Supplementary Fig. 6d**), suggesting that WWOX might limit SOX10 ubiquitination, although additional post-translational mechanisms cannot be excluded.

Collectively, these findings demonstrate that WWOX directly binds SOX10 via its WW1 domain, stabilizing SOX10 and reducing its proteasomal degradation. We propose a model in which WWOX preserves SOX10 function to support OPC differentiation and promote the maturation of myelinating oligodendrocytes.

## Discussion

Myelin is actively maintained throughout life to ensure axonal integrity and sustain the sensory-motor and cognitive functions of the nervous system. Efficient remyelination in response to demyelination is therefore essential for CNS homeostasis. Its loss can promote neurological symptoms and neurodegenerative diseases^10,11,13^. Here, using both human data and experimental models, we identify the WW domain–containing protein WWOX as a regulator of oligodendroglial differentiation and myelin repair. Our analyses of MS genetics and postmortem human tissue show that WWOX expression is associated with both disease risk and lesion dynamics. Developmental transcriptomics reveal that WWOX expression rises during gliogenesis, oligodendrogenesis, and active myelination, and remains enriched in OPCs in the adult brain. These observations motivated functional studies in vivo and ex vivo, through which we uncovered a previously unrecognized, cell-autonomous role for WWOX in oligodendroglial biology. Mechanistically, we identify SOX10, the master transcription factor for oligodendrocyte differentiation, as a direct binding partner of WWOX. This interaction stabilizes SOX10 and preserves its transcriptional activity during the generation of new myelinating oligodendrocytes.

While previous studies suggest that neuronal deletion of WWOX leads to hypomyelination via a non-cell autonomous mechanism^18,43^, our data demonstrate that oligodendroglia-specific deletion results in a distinct phenotype: OPC accumulation, reduced numbers of mature oligodendrocytes, and impaired remyelination following injury. Under baseline conditions, these defects are subtle, but when the system is challenged by cuprizone-induced demyelination, they become pronounced. WWOX-deficient OPCs mount the expected proliferative response to demyelination but fail to transition toward differentiation, remaining trapped in a premyelinating state. This arrest resembles the chronically stalled OPCs observed in non-repairing MS lesions. Notably, we find evidence of dysregulated WNT signaling, a pathway previously implicated in blocking OPC maturation^37^, suggesting a mechanistic basis for this differentiation deficit.

Evidence from rodent models indicate that remyelination failure is primarily mediated by new oligodendrocytes derived from OPCs. However, it could also be partially mediated by oligodendrocytes in demyelinated areas that regenerate their myelin sheaths and remyelinate demyelinated axons. Although our phenotypic characterization suggests a strong effect of WWOX deficiency in regulating the balance between OPC proliferation and differentiation, we also identified transcriptional alterations in oligodendrocyte cells and neurons. It is therefore possible that WWOX can also affect the remyelination potential of mature oligodendrocytes, and that non-cell autonomous processes may further exacerbate remyelination failure. Although our current study focuses on cell-autonomous mechanisms, remyelination failure might result from axonal damage perturbing cues required for myelination^10^. Future study of these non-cell autonomous processes will complement our findings. WWOX deficiency can therefore provide a valuable model to further explore the complementary of these signaling mechanisms.

Demyelination induced by cuprizone produces profound metabolic and inflammatory stress, driven in part by mitochondrial injury and oxidative damage in oligodendrocytes. WWOX is known to mitigate oxidative stress and preserve mitochondrial integrity^44–47^, and to limit inflammatory astrogliosis^43^. Our single-cell data showing a reduction in mature oligodendrocytes during demyelination in O-KO mice align with a model in which WWOX promotes both survival and stress tolerance in oligodendroglia. Thus, WWOX may act at multiple steps: ensuring OPCs can differentiate, sustaining oligodendrocytes under stress, and possibly shaping the glial environment needed for repair.

WWOX is a pleiotropic gene with established roles in cancer biology and neurodevelopment^18,48–50^. Our findings introduce an additional context in which WWOX contributes to human disease, myelin repair. Severe germline loss-of-function mutations in WWOX cause early developmental syndromes^50,51^, whereas somatic mutations can drive oncogenesis^52,53^. In contrast, MS risk is likely influenced by noncoding variants that subtly modulate gene expression. We propose that such regulatory variants could reduce WWOX levels just enough to weaken the WWOX–SOX10 interaction and destabilize SOX10, thereby diminishing the efficiency of remyelination. Although this model requires further testing, it suggests a promising direction: therapeutic strategies aimed at enhancing WWOX expression or stabilizing the WWOX–SOX10 interaction may boost myelin repair in MS and potentially other demyelinating disorders.

Taken together, our findings establish WWOX as a key regulator of oligodendroglial lineage progression and remyelination. By stabilizing SOX10 and enabling OPCs to proceed toward mature oligodendrocytes, WWOX emerges as a strong candidate for future remyelination-enhancing therapies.

## Supplementary Figures

**Supplementary Figure 1.**
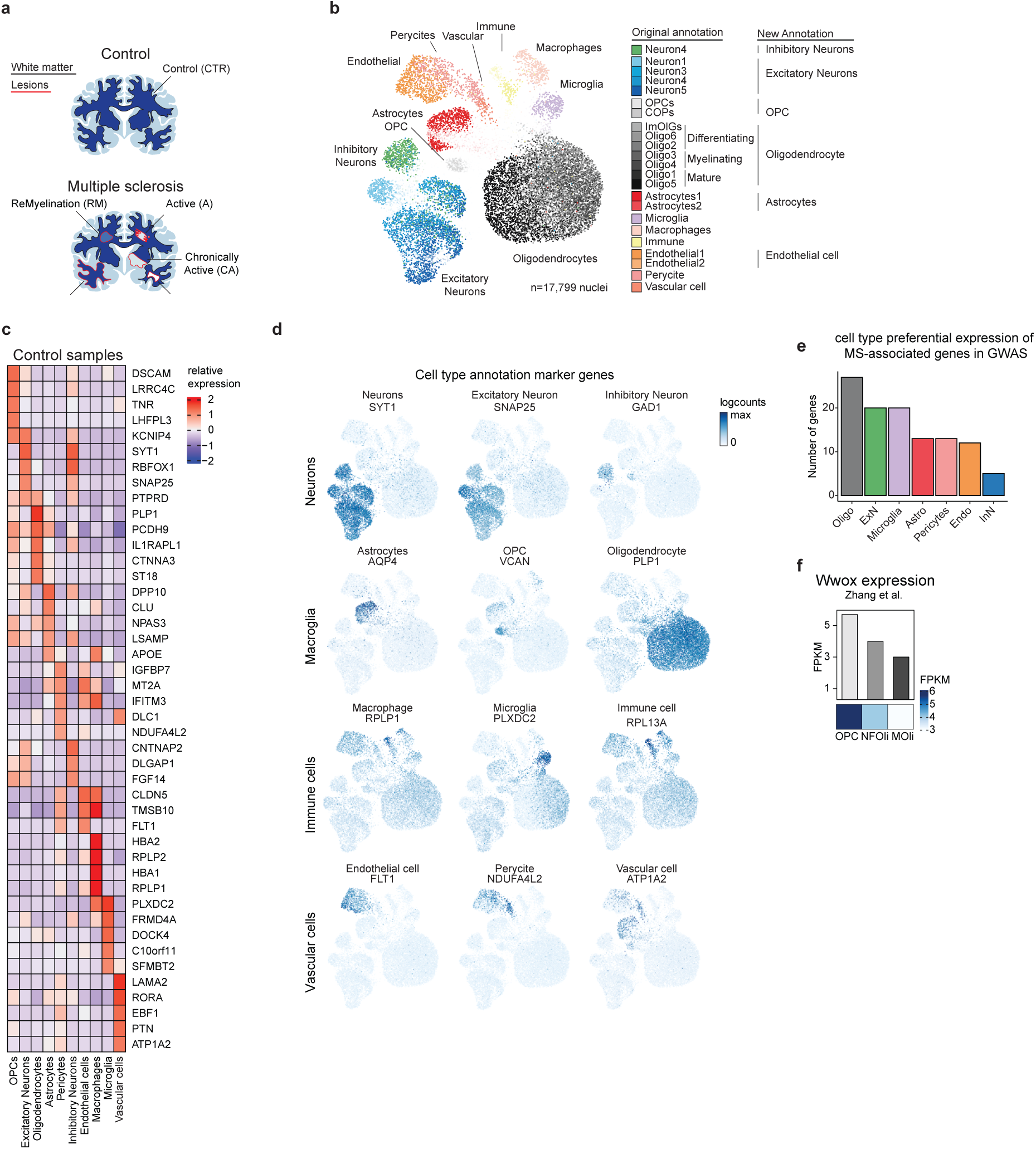
Annotation of MS and control post-mortem snRNA-seq data. **a** Schematics of control (CTR) and Multiple Sclerosis (MS) white matter lesion types. **b** 2D representation of nuclei from both CTR and MS postmortem samples. Each dot is a single nucleus and is labeled by the original annotation. Side annotations compares the original and our refined annotation. **c** Transcriptional signature expression of the top 5 signature genes for each cell type in the control condition. **d** Cell-type marker gene expression on the whole dataset (logcounts). **e** Number of cell-type-preferential genes prioritized in the original MS GWAS publication in each cell type. **f** Expression of Wwox in cell-sorted oligodendroglial bulk transcriptomics.

**Supplementary Figure 2.**
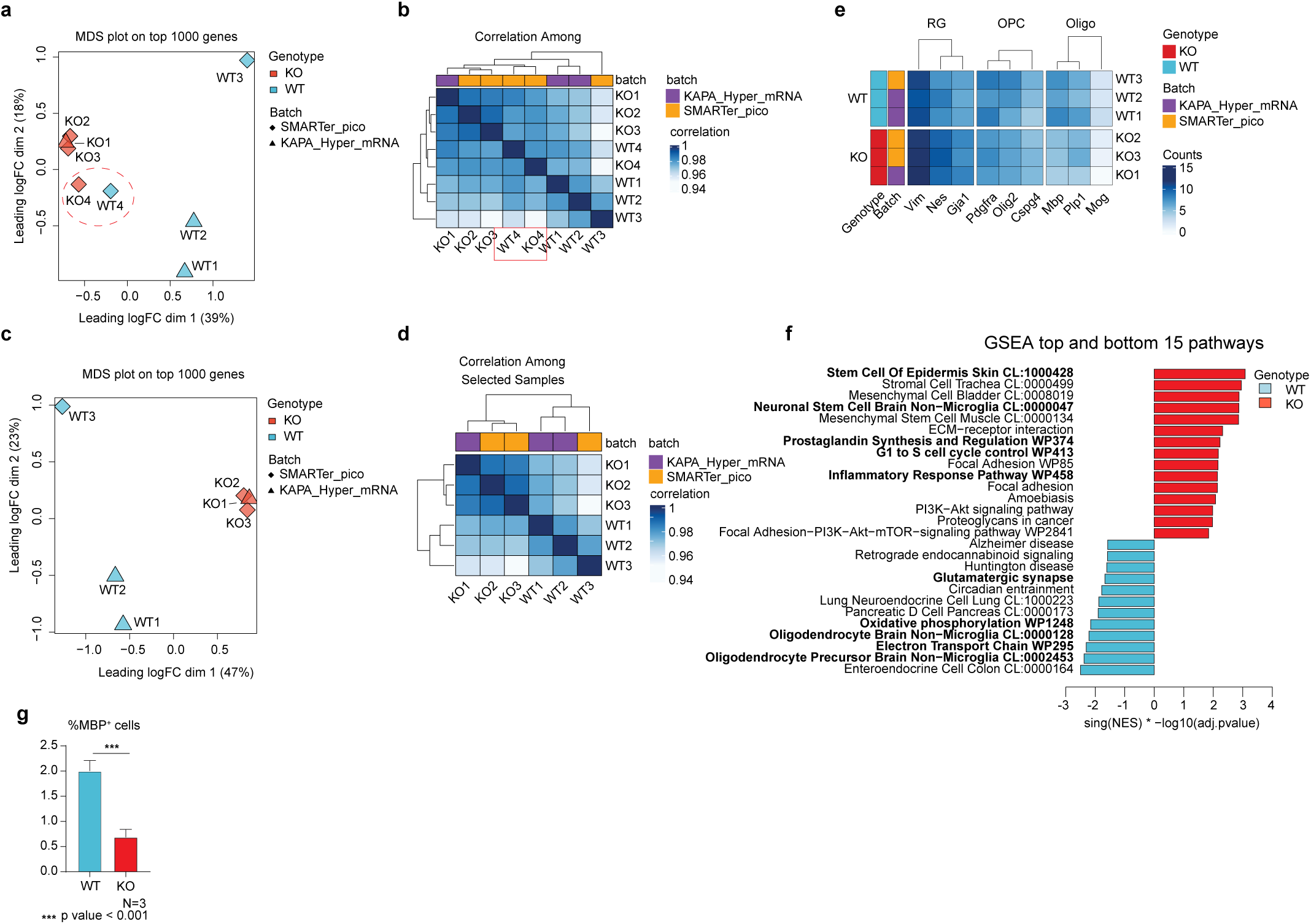
OPC culture quality controls and MBP quantification. **a** Multi-Dimensional Scaling (MDS) plot of the top 1000 genes for control (CTR) and KnockOut (KO) bulk OPC cell cultures. Low-quality replicates are highlighted(CTR4 and KO4). **b** Heatmap correlation of the whole transcriptome across all samples. Low-quality replicates are highlighted. **c-d** MDS and correlation heatmap after the removal of low-quality replicates. **e** Heatmap of cell-type marker genes expression across high-quality replicates. The expression is in corrected Counts Per Million (counts), obtained via ComBat_seq integration to corrected for sequencing kits as technical covariate, while using genotype as biological covariate. **f** Complete list of the most significant 5 over-and under-represented gene programs from each gene set(WikiPathway, KEGG, TabulaMuris). Selected pathways of Fig.2 are highlighted in bold. **g** Quantification of Olis (MBP+) in OPC culture after 7 days in vitro from the differentiation (related to Fig. 2e).

**Supplementary Figure 3.**
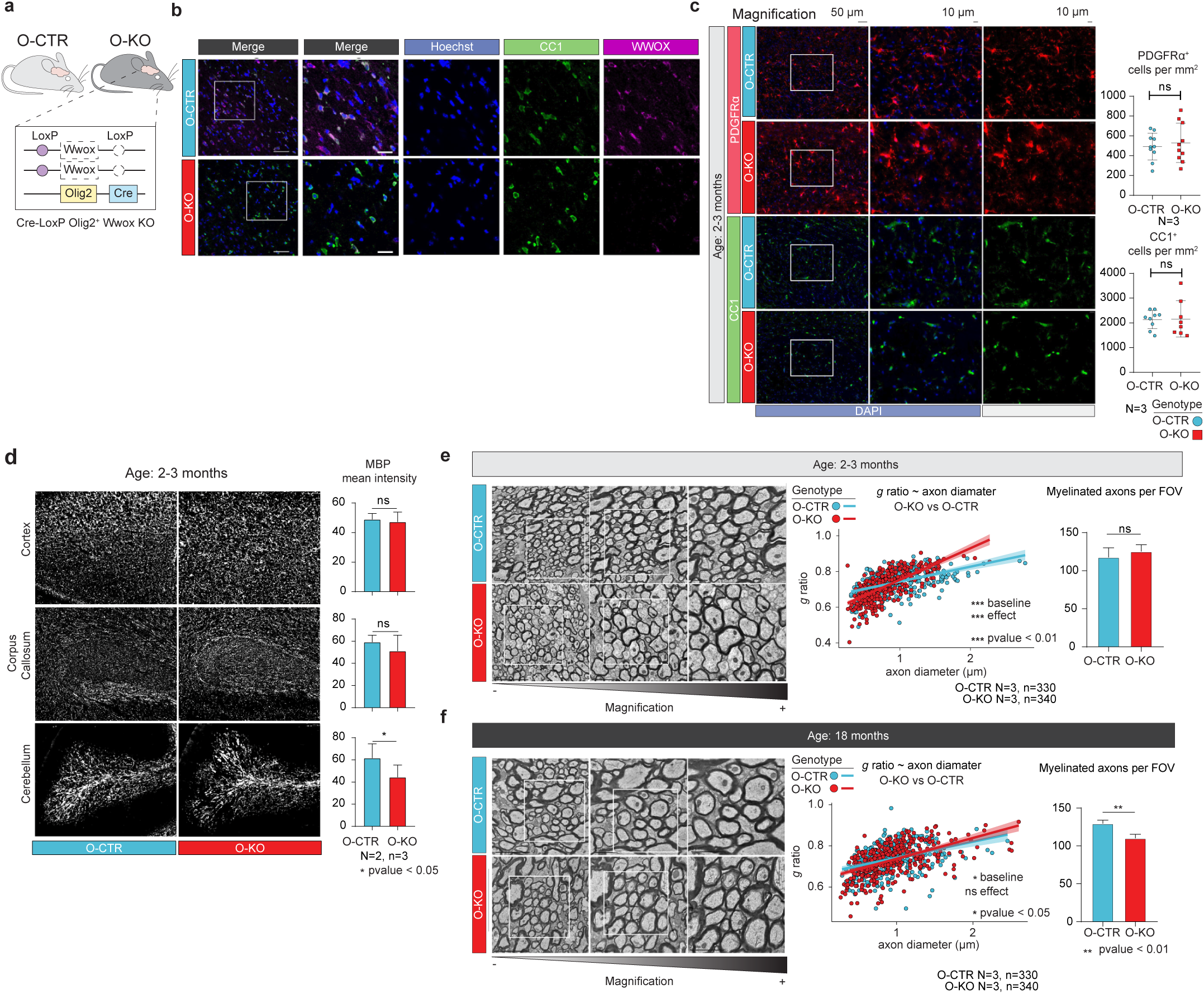
Conditional deletion of WWOX in OPCs affects remyelination in ageing mice. **a** Schematic of the Cre-LoxP-based genetic mouse model used to conditionally delete *Wwox* in oligodendrocyte lineage under Olig2-Cre expression. **b** IF staining for WWOX and CC1 (mature Oli) in the corpus callosum of O-CTR and O-KO mice at Postnatal day 17 (P17). WWOX expression is markedly reduced in CC1^+^ Olis of O-KO brains, confirming efficient Cre-mediated recombination. **c** IF staining and quantification of OPCs (PDGFRα+) and Olis (CC1+) in the corpus callosum (data are shown as means ± SEM; two-tailed t-test). **d** IF for MBP in O-CTR and O-KO brain regions at 2 months of age. Quantification was based on N=2 mice per genotype, with n=3 sections analyzed per animal (data are shown as means ± SEM; two-tailed t-test). **e** Representative TEM images of the midsagittal corpus callosum from adult O-CTR and O-KO mice. White squares indicate magnified regions. Middle: Quantification of myelin thickness by g-ratio analysis (linear regression). Right: Quantification of the number of myelinated axons per FOV shows no significant difference between groups (data are shown as means ± SEM; two-tailed t-test). **f** Repeated analysis shown in **e** with 18-month-old O-CTR and O-KO mice.

**Supplementary Figure 4.**
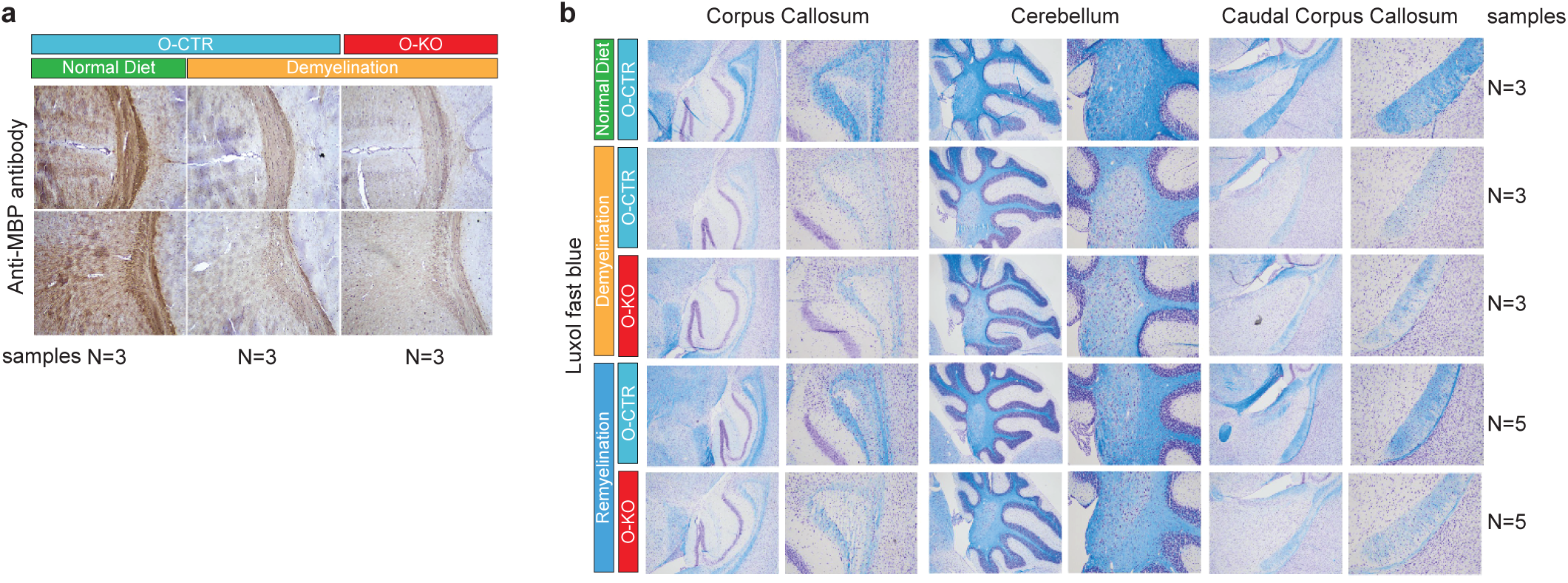
Assessment of demyelination and remyelination in O-CTR and KO mice. **a** Representative images of coronal brain sections stained with anti-MBP antibody for O-CTR and O-KO following cuprizone-induced demyelination (data are shown as means ± SEM; two-tailed t-test). **b** Luxol Fast Blue (LFB) staining of multiple brain sections across conditions. Paraffin-embedded brain sections were stained with LFB, where myelin fibers appear blue and neuronal cell bodies violet. Reduced myelin staining was observed in both O-CTR and O-KO mice following demyelination relative to ND controls. Following 2 weeks of remyelination, reduced blue staining remained evident in O-KO mice compared to O-CTR across all three brain regions.

**Supplementary Figure 5.**
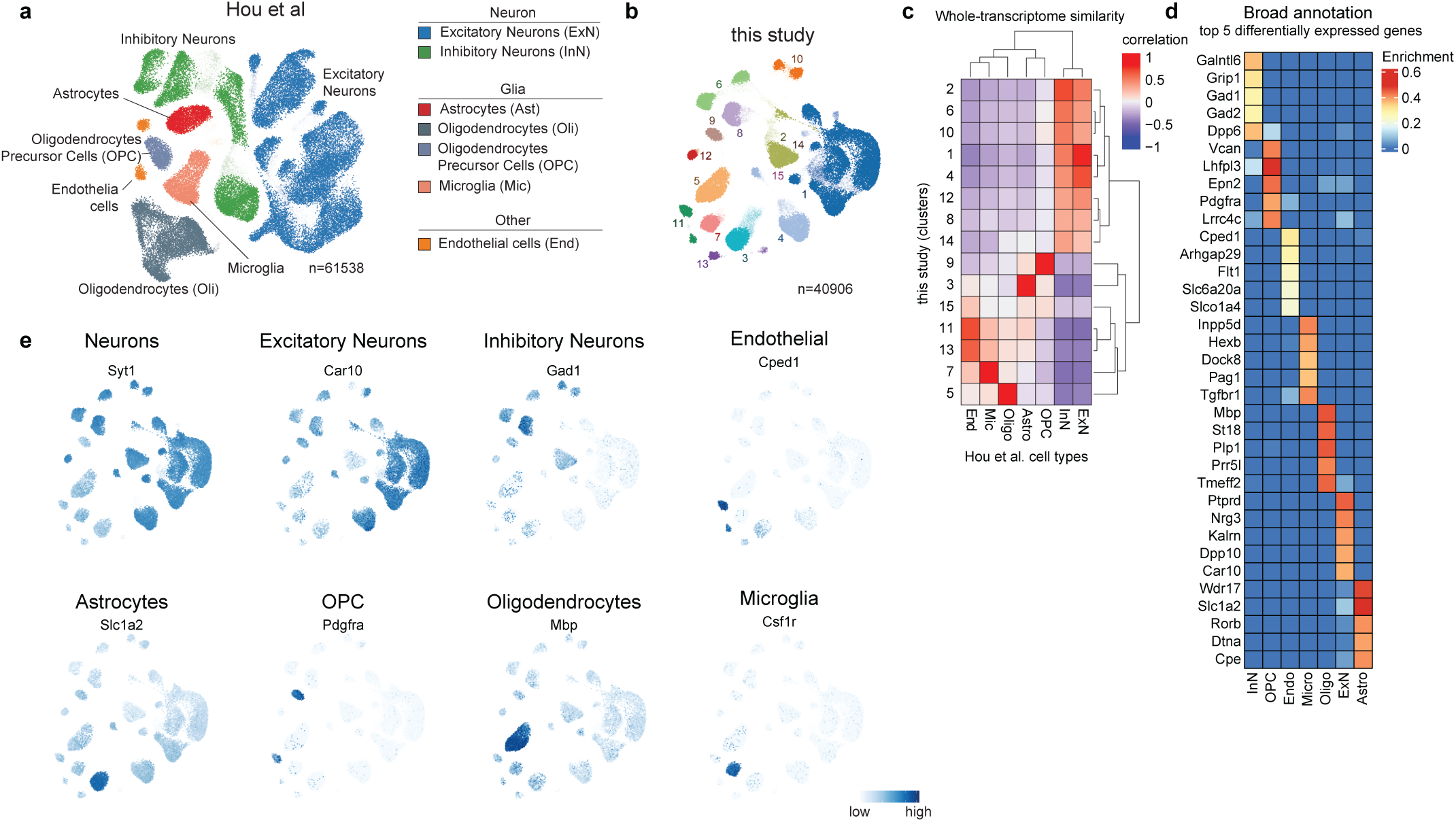
Annotation of O-CTR and O-KO snRNA-seq data across diet conditions. **a** 2D representation of Hou et al. 61,538 nuclei, labelled by cell type. Hou et al. snRNA-seq data were used as a reference for cell-type annotation. **b** Communities identified in our snRNA-seq data via Leiden clustering and used for cell-type annotation. **c** Correlation heatmap of transcriptional signatures between Hou et al. cell types and cluster communities. Hierarchical clustering separates neuronal and glial populations. **d** Heatmap of the top 5 enriched genes per cell type. Enrichment scores were obtained via a hypergeometric enrichment test. **e** Cell-type marker gene expression visualized on the 2D representation of the data (expression is in logcounts).

**Supplementary Figure 6.**
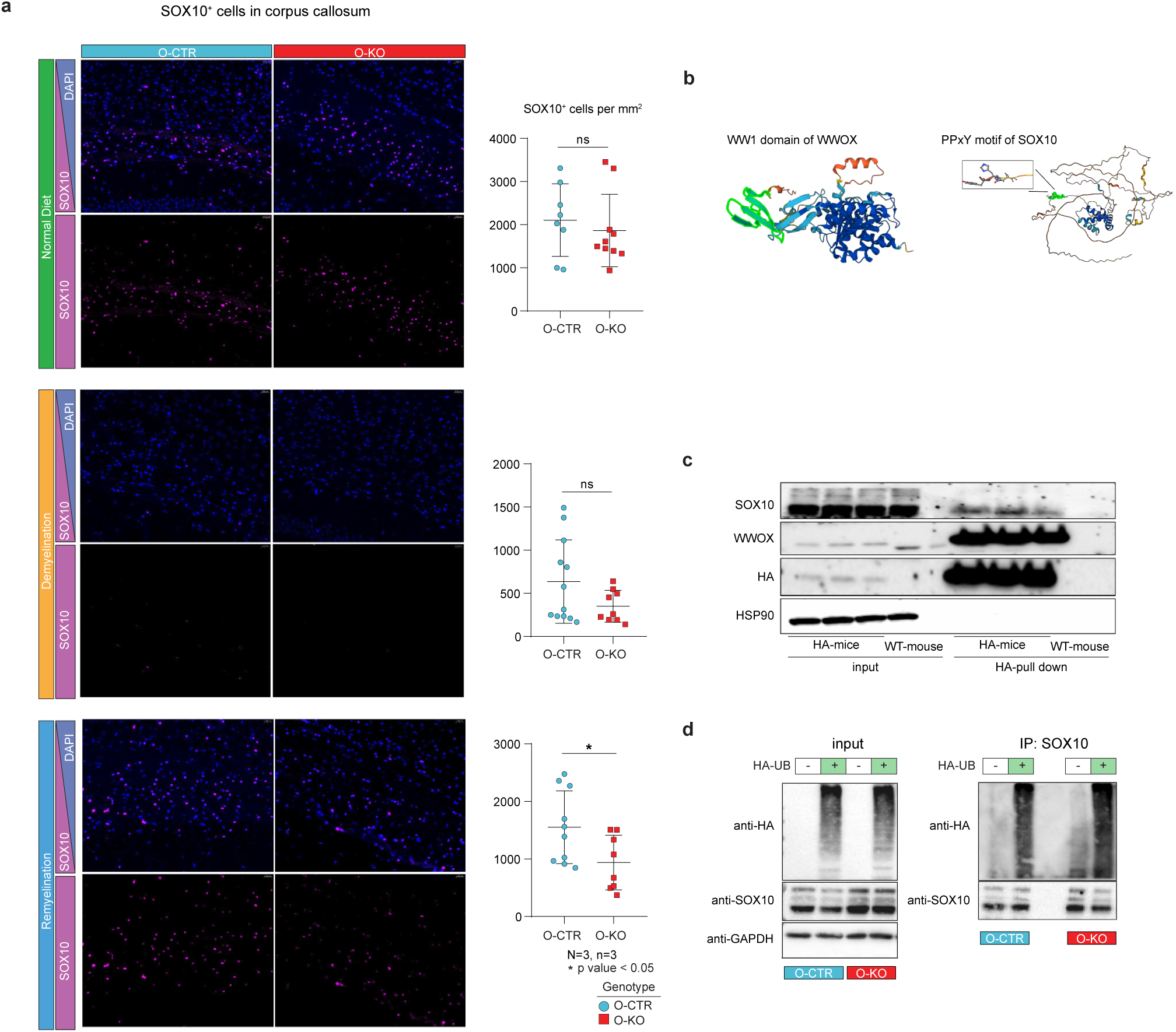
The physical interaction between WWOX and SOX10. **a** IF staining and quantification of SOX10+ cells in the corpus callosum of O-CTR and O-KO mice across conditions (normal diet, demyelination, and remyelination).(data are presented as mean ± SEM; two-tailed t-test; n.s., not significant). **b** AlphaFold structural predictions of WWOX and SOX10 proteins. The WW1 domain in WWOX and the PPxY motif in SOX10 are highlighted, suggesting a potential interaction interface. **c** Western blot of HA pull-down assay using cortical lysates from HA-tagged WWOX mice. **d** Western blot analysis of WT and Wwox-KO Oli-neu cells transfected with HA-Ubiquitin (Addgene plasmid #18712). Lysates were subjected to immunoprecipitation using anti-SOX10 antibody followed by immunoblotting with anti-HA to assess ubiquitination of SOX10.

## Methods

### Data sources

#### Reference single-cell transcriptomics data

snRNA-seq data of the white matter of post-mortem tissue of individuals with progressive MS and human controls (CTR) was obtained from^15^. Reference mouse snRNA-seq data for cell annotation via signature-based label transfer was obtained from^34^.

#### Reference bulk transcriptomics data

developmental forebrain gene expression data across the human lifespan were retrieved from^27^. Expression trends were analyzed by applying smoothing Reads Per Kilobase Million (RPKM) values using the geom_smooth function with loess regression and 0.25 span in ggplot2. Multiple span values were tested to avoid artefacts due to the smoothing of expressions across discrete time points. A vertical line was added to highlight birth time.

#### GWAS enrichment analysis

The association between cell-type-specific genes and MS genetic scores was tested using MAGMA^54^. Cell-type-specific gene scores were estimated considering only control samples, and cell types were compared one against all the others by performing differential expression analysis using NEBULA^55^. Briefly, NEBULA fits a negative binomial linear mixed model to account for cell-subject membership associations. For comparative analyses, cells of the same lineage were grouped together, except neuronal subtypes (inhibitory and excitatory neurons). Only protein-coding genes detected in at least 10% of cells were tested, and Benjamini-Hochberg correction was applied to adjust p-values. A cell-type preferential expression score for each gene was computed as −log10(adjusted p-values) * log2(FC). Gene-level GWAS association scores were computed using MAGMA as previously described in^22^. Associations between preferential expression and gene-level genetic risk scores were tested with MAGMA gene-level analysis with a continuous covariate for expression. Candidate genes driving cell-type specificity and GWAS associations were identified as preferentially expressed for a candidate cell type (p-value < 0.01 NEBULA analysis), prioritised by the International Multiple Sclerosis Genetics Consortium^23^, and differentially expressed in MS vs CTR (see below). Genes were ranked by gene-level GWAS association score. Preferential gene expression was quantified as log2(FoldChange) of candidate cell type vs other cell types; the genetic score as −log10(adjusted p-value); MS differential expression as −log10(adjusted p-value). Differential expression of oligodendroglia genes in each MS lesion or CTR condition was tested versus all other cells using the Wilcoxon signed-rank test. A gene was considered differentially expressed if its absolute log(FoldChange) was higher than 0.05, was detected in at least 25% of the cells, and had an adjusted p-value lower than 0.01.

### Cell culture

#### Primary OPC isolation and culture

Mixed glial cultures and OPCs were isolated using the differential attachment method. Briefly, brains from P0–P2 mouse pups were isolated and placed in ice-cold HBSS. Meninges were carefully removed under a dissection microscope. Brain tissues were enzymatically dissociated using 0.01% trypsin and 10 μg/ml DNase at 37 °C for 7–10 minutes. Following enzymatic digestion, the cells were washed with ice-cold DMEM, passed through a 70 μm cell strainer, and plated onto poly-D-lysine-coated T-25 Flasks.Cells were cultured to confluence (DIV 7-10) in a humidified incubator at 37 °C with 5% CO_2_ in mixed glia medium composed of DMEM/F-12 supplemented with 10% FBS and 1% Penicillin-Streptomycin (Pen/Strep). For OPC isolation, the flasks were shaken at 37 °C for 30 minutes at 140 rpm to remove microglia, followed by overnight shaking at 220 rpm. The resulting cell suspension was plated onto uncoated petri dishes to allow further removal of microglia and astrocytes by differential adhesion. Highly enriched OPCs were collected by centrifugation and resuspended in serum-free OPC growth medium consisting of DMEM supplemented with 1% Pen/Strep, 1X B27, 1% sodium pyruvate, 1% glutamate, and 1% Horse-serum. For differentiation, 340 ng/ml T3 (triiodothyronine) and 400 ng/ml T4 (L-thyroxine) were added to the culture medium.

#### Oli-neu cell line maintenance and Wwox knockout generation

The mouse oligodendrocyte progenitor cell line, Oli-neu, were a generous gift from the laboratory of Prof. Elior Peles, Department of Molecular Cell Biology, Weizmann Institute of Science. To knock out *Wwox*, lentiviral particles were generated using the LentiCRISPR-v2 plasmid (Addgene plasmid #52961) encoding an sgRNA targeting exon 4 of *Wwox* (5′-CACCGCACCAGACAGAGATACGAC-3′). Oli-neu cells were transduced with the viral supernatant and subsequently selected with 1 μg/mL puromycin to enrich for infected cells (KO). Non-transduced Oli-neu cells were used as controls (WT). Knockout efficiency was confirmed by Western blot analysis. Oli-neu cells were maintained in an undifferentiated state in DMEM supplemented with 1% penicillin–streptomycin, 1% glutamine, 2% B27, 10 μg/mL insulin, 1% sodium pyruvate, 20 ng/mL PDGF-AA, 1 ng/mL neurotrophin-3 (NT-3), 10 ng/mL ciliary neurotrophic factor (CNTF), and 4.2 ng/mL forskolin.

### Animal models

#### Ethical approval

All mouse procedures were performed and approved by the Hebrew University Institutional Animal Care and Use Committee (HU-IACUC).

#### Wwox ablation

Mice carrying two loxp sites *(Wwox ^flx/flx^)* adjoining exon 1, previously established in our lab Abdeen, S. K. et al.(2013), were used to conditionally ablate *Wwox* in OPCs. *Wwox* conditional ablation from OPCs was achieved through crossing *Wwox ^flox/flox^* mice with Olig2-Cre mice purchased from Jackson Laboratory (Stock: 025567, B6.129-Olig2tm.11(cre)Wdr/J). *Wwox ^flx/flx^* mice harbouring a Cre-recombinase have a conditional homozygous *Wwox* deletion, the *Wwox ^flx/flx^*; *Olig2 ^cre/+^* mice are referred to as O-KO. Conditional *Wwox* knockout mice further had a reporter allele, the Rosa26-loxp-STOP-tdTomato. All the conditional *Wwox* knockout mice were maintained in a mixed genetic background of C57BL6/J;129sv. *Wwox* WT (*Wwox ^+/+^*,WT), heterozygous (*Wwox ^+/-^*, Het and null (*Wwox ^-/-^,* KO) mice were previously generated in our lab and were all maintained on an FVB background R.I. Aqeilan (2007). All mice were maintained in a specific pathogen-free (SPF) unit with access to food and water and a 12-hour light/dark cycle.

Genotyping was performed on DNA extracted from a tail or earpiece of weaning mice using DNA lysis buffer prepared with autoclaved double-distilled water (DDW), 10 mM sodium hydroxide (NaOH) and 300 µM EDTA. PCR reaction was further performed using the following primers:

1. *Conventional WT:* Forward primer 5’ to 3’: “GCAGAATGTCTTGCTAGAGCTTTG”, Reverse primer 3’ to 5’: “ATACTGACATCTGCCTCTAC”
2. *Conventional KO*: Forward primer 5’ to 3’: “GCAGAATGTCTTGCTAGAGCTTTG”, Reverse primer 3’ to 5’: “CAAAAGGGTCTTTGAGCACCAGAG”
3. *Rosa WT:* Forward primer 5’ to 3’: “AAGGGAGCTGCAGTGGAGTA”, Reverse primer 3’ to 5’: “CCGAAAATCTGTGGGAAGTC”
4. Rosa Tomato: Forward primer 5’ to 3’: “CTGTTCCTGTACGGCATG”, Reverse primer 3’ to 5’: “GGCATTAAAGCAGCGTATCC”
5. *Olig2-Cre:* Forward primer 5’ to 3’: ”GCATCATTCGAGAGGTTAGA“, Reverse primer 3’ to 5’: “TTACGGCGCTAAGGATGACT“, Olig2-cre Common: “CTTTCTTGGTGGAAGACGTG”

#### Cuprizone-induced demyelination and remyelination

Six- to eight-week-old mice were fed a 0.2% cuprizone-containing diet (Envigo, Catalog No. TD.140803) for 5 weeks to induce demyelination. To assess remyelination, mice were returned to a normal diet for 2 weeks. Age-matched control mice were maintained on a normal diet throughout the experiment. Cuprizone pellets were stored in vacuum-sealed packages, protected from light, at 4 °C until use. The chow was replaced every other day. Mice were monitored daily for food intake and three times per week for body weight. Brain tissues were collected at two time points: at the end of the 5-week cuprizone treatment to assess demyelination, and 2 weeks after reversion to a normal diet to evaluate remyelination.

### Immunofluorescence staining

Mice were euthanised using CO_2_ and transcardially perfused with 4% paraformaldehyde (PFA) in PBS. Dissected brains were post-fixed overnight at 4 °C, then cryoprotected in 30% sucrose at 4 °C overnight. Brains were embedded in OCT compound, and sagittal sections (12–14 μm) were prepared using a cryostat. Sections were washed with PBS, permeabilised in 0.1% Triton™ X-100 in PBS (PBT), and blocked with 5% goat serum containing 0.5% Triton™ X-100 for 1 hour at room temperature. Subsequently, sections were incubated with primary antibodies overnight at 4 °C. After washing with PBS, sections were incubated with Alexa Fluor-conjugated secondary antibodies for 1 hour at room temperature. Finally, sections were washed and mounted using DAKO mounting medium (Catalog No. S3023).

List of the antibodies used and their working dilution:

| Antibody | Dilution | Catalog No. |
| --- | --- | --- |
| Rabbit Anti-MBP | 1:200 | Ab65988 |
| Mouse Anti-CC1 | 1:50 | OP80 |
| Rabbit Anti-WWOX | 1:10,000 | N.A. |
| Rabbit Anti-SOX10 | 1:200 | ab227680 |
| Rat Anti-PDGFRα | 1:500 | 558774 |
| Goat Anti-Mouse IgG, Alexa flour 488 | 1:1000 | Ab150105 |
| Goat Anti-Rabbit IgG, Alexa flour 647 | 1:1000 | ab150079 |
| Goat Anti-Rabbit IgG, Alexa flour 488 | 1:1000 | ab150077 |
| Alexa Fluor 488-conjugated donkey anti-Rat IgG | 1:500 | 712-545-153 Jakson |
| Donkey Anti-Rat IgG, conjugated with Cy5 | 1:500 | 712-175-150 |

#### Image acquisition and analysis

Immunofluorescence-stained sections were imaged using an Olympus FV1000 confocal microscope and 3 DHISTECH panoramic scan. The obtained images were processed using the F-10-ASW viewer and slide viewer. Images were further analysed using ImageJ and QuPath-0.5.1. Sholl analysis and cellular surface area measurements were performed using ImageJ Sholl plugin.

#### Statistical analysis

Quantifications for immunofluorescence stainings and electron microscopies images were presented using GraphPad Prism 8. Results are shown as mean ± standard error of the mean (SEM). P values were computed using Student’s t-test, one-sample t-test, Kolmogorov-Smirnov, Mann-Whitney test, repeated measures analysis of variance (ANOVA), and Spearman’s correlation test, with p value < 0.005 being considered significant (* p value < 0.05, ** p value < 0.01, *** p value < 0.001).

### Bulk RNA-sequencing of oligodendroglia from mouse primary cultures

#### Sequencing and data preprocessing

Bulk RNA-seq data were generated for 48-hours differentiated primary OPC cultures with KAPA mRNA HyperPrep Kits or SMARTer pico Kits, depending on the available starting material. After sequencing, quality control were applied to reads files, including trimming of low-quality bases, residual adapters and polyA tails, via Trim_Galore v0.6.1 (Krueger, n.d.) and FastQC (“Babraham Bioinformatics - FastQC A Quality Control Tool for High Throughput Sequence Data,” n.d.). Short reads (<20bp) were filtered out. Subsequently, reads were aligned to the transcriptome GRCm39 using Salmon^56^. Only protein-coding genes with at least one detected gene were retained for the analysis, resulting in a total of 19,777 genes. Sequencing artefacts associated with different sequencing kits were removed via batch correction using ComBat_seq^57^. Sequencing kits were specified as batches, while conditions (WT and KO) were used as groups to preserve biological differences. Corrected counts were normalised as Trimmed Mean of M values (TMM) using edgeR^58^ and transformed as log2(TMM + 1) values.

#### Differential expression analysis

Differential Expression Analysis (DEA) was performed with edgeR. Briefly, genes were filtered to remove lowly expressed genes, using filterByExpr with default parameters. TMM normalisation was performed again to account for removed genes, and gene-wise dispersion was estimated using estimateDisp, with default parameters and conditions (WT and KO) specified in the design matrix. A negative binomial generalised log-linear model was fitted using glmQLFit, and a gene-wise quasi-F-test was performed to assess differential expression via glmQLF. To account for the high number of tests, P-values were adjusted using the Benjamini-Hochberg (BH) correction with the topTags function. A gene was considered differentially expressed if its log2(Fold Change) was higher than 1 and an adjusted p-value lower than 0.05.

#### Gene set enrichment and variational analysis

Gene Set Enrichment Analysis (GSEA) was performed using fgsea (https://github.com/alserglab/fgsea) to assess statistical over- or under-representation. Scores for GSEA were defined as *sign(log2(Fold Change)) * −log10(adjusted pvalue)*. Tabula Muris, WikiPathways, and KEGG gene sets collections were retrieved from Enricher (https://maayanlab.cloud/Enrichr/). Adjusted p-values were computed via the BH correction. Gene sets with an adjusted p-value lower than 0.01 were retained and split into two groups: overrepresented in Wwox-depleted, with a positive Normalised Enrichment Score (NES), and underrepresented with a negative NES, for each of the three sets. For each set and split, gene sets were ranked by increasing adjusted p-value. Top-thirty enriched gene sets were selected for gene set variational analysis. Gene set activity scores per sample were estimated by applying Gene Set Variational Analysis (GSVA) ^59^ on log2(TMM + 1) values.

### Electron microscopy

#### Tissue processing and quantification

Mice at different time points were euthanised with CO_2_ and perfused transcardially with a fixative solution composed of 2% PFA, 2.5% glutaraldehyde in 0.1 mM sodium cacodylate buffer with pH equivalent to 7.3. Mice brains were dissected, placed in the fixative solution for another 2 hours at RT and stored afterwards at 4° C till the time of processing. Brains were further cut using an altro-acrylic 1mm brain matrix (Bar Naor, #BS-A-5000C) to collect the corpus callosum (CC). The collected CC was washed with sodium cacodylate, fixed with another fixative solution consisting of 1% osmium tetroxide, 1.5% potassium ferricyanide in sodium cacodylate and then washed four times again with the same fixative solution. Samples were dehydrated with 30, 50, 70, 80, 90 and 95% EtOH gradually for 10 min each and then placed in 100% EtOH for three times 20 min each. Samples were then put in two changes of propylene oxide. Afterwards, they were infiltrated 25,50,75 and 100% epoxy resin gradually for 24 hours per each one and then placed in the oven at 60° C for 48 hours to make the tissue blocks. The blocks were further sectioned to 80 nm sections using an ultramicrotome and stained with lead citrate and uranyl acetate. Stained sections were then viewed using a transmission electron microscope (TEM) of Joel JEM 1400 Plus and EM micrographs were captured by a Gatan Orius CCD camera, and analysed using ImageJ. For g-ratio calculations, the inner axonal diameter of 300 myelinated axons, 100 axons per mouse, was measured and divided by the outer total myelinated axonal diameter.

#### g ratio ~ axon diameter quantification

Linear models were fitted using lm from stats (Ripley 2001) to predict the g ratio using an interaction term between the axon diameter and genotype. Significance for the intercepts, representing the base amount of myelin, and the interactions term are reported. Linear model assumptions were tested for each model. Linearity of the data was tested using the Kolmogorov-Smirnov test. Zero mean of the residuals was assessed by checking that the 25th and 75th percentiles fall between 0±0.1 interval from the mean. Autocorrelation of residuals and constant residual variance were tested using the Durbin-Watson test and non-constant residual variance test, respectively, via car (Fox and Weisberg 2018). Results for the test were considered significant for a p-value lower than 0.05.

### Single-nucleus RNA-sequencing of mouse cortex

To investigate the transcriptional changes associated with *Wwox* deletion in oligodendrocyte precursor cells (OPCs), single-nucleus RNA sequencing (snRNA-seq) was performed on cortex samples from *Olig2-Cre Wwox KnockOut (O-KO)* and *Wild-Type (O-WT)* mice, with balanced sex proportions (half male, half female). Samples were collected under three experimental conditions and replicates were pooled: (1) normal diet, two replicates per genotype; (2) cuprizone-induced demyelination, two replicates per genotype; (3) remyelination following cuprizone withdrawal, three replicates per genotype.

#### Nuclei isolation

Nuclei were isolated from frozen cortical and white matter tissue using a previously published protocol with minor modifications^34^. Briefly, tissue was Dounce homogenised in 5 mL of lysis buffer containing 10 mM Tris-HCl (pH 7.4), 10 mM NaCl, 3 mM MgCl_2_, and 0.025% NP-40 for 15 minutes. The homogenate was filtered through a 30 μm cell strainer and centrifuged at 500 × g for 5 minutes at 4 °C. The nuclei pellet was washed and filtered twice using nuclei wash buffer (1% BSA in PBS with 0.2 U/μL RNasin, Promega). Nuclei were resuspended in nuclei wash buffer at a concentration of 1,200 nuclei/μL.

#### Library preparation and sequencing

Isolated nuclei were loaded onto a Chromium Next GEM Single Cell Gene Expression v3.1 (3’) kit (10x Genomics), and libraries were prepared according to the manufacturer’s protocol. The scRNA-seq libraries were sequenced on the Illumina Novaseq6000 sequencing system in a 28-bp and 90-bp as paired-ended. Fastq files were processed using CellRanger v7.2.0 with mm10-2020-A genome to estimate the filtered count matrices.

#### Preliminary quality controls

Raw counts of 51,066 nuclei were filtered based on standard quality metrics. Nuclei with fewer than 500 (empty droplets) or more than 10000 genes detected (multiplets) were filtered out. Ensembl IDs were converted into Gene IDs, and gene biotypes were retrieved. Only protein-coding genes were retained, and genes detected in less than 10 nuclei were removed. High-mitochondrial nuclei were identified via K-means clustering using the ratio of mitochondrial vs non-mitochondrial reads and searching for two groups considered high-mitochondrial and high-quality nuclei. After the first step of quality controls, 47,960 nuclei and 17,326 genes were retained for further quality controls.

#### snRNA-seq data analysis

All dimensionality reduction, clustering, and vidualization analyses were performed using the ACTIONet framework^60^ using default parameters. For cluster annotation, cluster signatures were computed and correlated with the pre-annotated reference mouse data^34^ using Pearson’s correlation. Each cluster was annotated using the most correlated cell-type label. Seven major communities were identified: excitatory neurons, inhibitory neurons, astrocytes, oligodendrocyte precursor cells, oligodendrocytes, microglia and vascular cells. The ACTIONet framework was reapplied with default parameters for each major community to identify sub-communities. Multiple quality metrics (total number of counts, number of mitochondrial counts, and cell-type marker gene expression) were controlled for each sub-community, marking out-of-distribution communities as low-quality cells. To detect doublets, scds^61^ was applied to the whole data set to compute single nuclei doublet scores. Cells were classified in two groups by applying K-means clustering on doublet scores. After removing low-quality nuclei and doublets, 40,906 high-quality nuclei and 17,326 genes were retained for further analysis. Nuclei annotation was improved by performing Leiden clustering on the clean data using a resolution of 0.1, obtaining coarse communities for transcriptional signatures. Resolutions between 0.1 and 2 were tested to select the best community resolution for reference-based annotation. Signatures-based label transfer was repeated to annotate high-quality communities (min: 0.18, median: 0.68, max: 0.80 correlation). Seven broad high-quality cell types were identified: 18911 excitatory and 9483 inhibitory neurons, 3722 oligodendrocytes, 4217 astrocytes, 2022 microglia, 1077 oligodendrocyte precursors and 1474 endothelial cells. ACTIONet was used to visualise marker gene expression on the UMAP projection of single nuclei. Logcounts of selected marker genes were visualised after imputation of gene expression using the function *imputeGenes* with default parameters.

#### Simulated replicates and abundance analysis

Differences in proportions were tested with three simulated replicates for each combination of genotype and condition. Simulated replicates were obtained with an unbalanced sampling of 33% of the nuclei without replacement. Abundance shifts between genotypes were tested using MiloR^35^. Briefly, ACTION compressed representation of the data was obtained with ACTONet with *reduce.ace*. All 50 dimensions of the ACTION representation were used to build the KNN network, with 50 nearest neighbours for all the cells. Multiple tests with different numbers of neighbours were performed to achieve the community dimensions suggested in MiloR. Fine-grained communities were defined for representative cells via refined sparse sampling. For each community, differences in genotype abundance were assessed using a negative binomial generalised linear model. To account for shared cells between communities, spatial false discovery rate was used to correct p-values. Annotation of communities with cell-type labels was performed using *annotateNhoods* with default parameters. Abundance results were visualised in a Bee Swarm plot using *plotDAbeeswarm* for each condition.

#### Mouse snRNA-seq differential expression analysis

DEA was performed with the Wilcoxon-Mann-Whitney test implemented in presto (https://github.com/immunogenomics/presto) comparing control and Wwox-depleted samples across matched cell types and conditions. Adjusted p-values were computed using the BH correction. A gene was considered differentially expressed for abs(logFC) > 0.25 and an adjusted p-value < 0.05. Multiple thresholds of abs(logFC) were tested (0.05, 0.25, 0.5) to ensure the overall robustness of the results.

#### Oligodendroglia differential trajectory inference

The condiments^36^ framework was used to test trajectory differences between genotypes. Before estimating trajectories, intermediate communities of the oligodendroglia subset (OPC, Committed Oligodendrocytes, Oligodendrocytes and Damaged Oligodendrocytes) were identified via Leiden clustering at 0.3 resolution. Such communities were validated by visually inspecting marker genes for oligodendroglia differentiation retrieved from literature. Remyelinating oligodendroglia was isolated and re-analysed with ACTIONet for each condition, obtaining a specific ACTION compressed representation and UMAP visualisation. Trajectories across intermediate communities were computed via slingshots^62^ on all 50 dimensions of the ACTION compressed representation. The OPC community was set as the starting point. Differences in trajectories between genotypes were tested using topologyTest from condiments with 100 replicates. If no differences in topology were found, differences in progression could be tested via progressionTest with default parameters. For both tests, differences were considered significant for p-values lower than 0.01. Trajectory estimation with slingshot was repeated on the 2D coordinates of the UMAP only to visualise trajectories. A minimum spanning tree was obtained using *slingMST* and visualised on the UMAP projection using ggplot2.

#### Gene set enrichment analysis

Gene Set Enrichment Analysis (GSEA) was performed using fgsea. Condition-specific scores were computed for OPCs and oligodendrocytes from the DEA summary statistics:sign(log2(Fold Change)) −log10(adjusted pvalue). Mouse-specific WikiPathways, KEGG, and REACTOME gene sets were obtained from MsigDB using the msigdbr package. Significant gene sets (adjusted p-value < 0.05) were ranked based on NES for each cell type and condition combination. GSEA was performed to test the over- or under-representation of TFs’ putative targets. A-level positive transcription factor regulons for SOX10 were obtained from Dorothea^41^. Gene sets were considered significant with an adjusted p-value lower than 0.05.

#### Cell cycle estimation

Cell-cycle scores for OPCs were estimated using tricycle^38^. Nuclei were projected on the tricycle’s cell-cycle mouse brain reference, with shared 499 genes. Cell-cycle scores were estimated using *estimate_cycle_position* with default parameters. Top2a and Smc2 expression peaks were detected and centred on π, suggesting good-quality cell-cycle scores. Genotype differences in OPC cell-cycle score distributions were tested via the Anderson-Darling k-Sample test for each condition using the kSamples package. Density plots of OPCs were obtained using *geom_density* with ggplot2.

#### Pseudobulk estimation

Pseudobulk samples were computed for each cell type-genotype-condition combination by a gene-wise sum of total counts and normalised as log2(CPM + 1).

### WWOX-SOX10 interaction test via GST pull-down assay

#### GST pull-down assay

HEK293T cells were plated in appropriate culture plates and allowed to adhere overnight. The following day, cells were transfected with a *SOX10*-inducible plasmid (addgene #115241) along with a *GST-tagged WWOX* plasmid^63^. For transfection in 6-well plates, 0.3 μg of *SOX10* plasmid was combined with 0.1–0.3 μg of *GST-WWOX* plasmid. Lipofectamine reagent was used at a ratio of 3 μL per 1 μg of DNA, diluted in 100 μL of serum-free medium (SFM) per 1 μg of DNA. Cells were incubated with the transfection mixture for 48 hours. To induce SOX10 expression, doxycycline (DOX) was added at a final concentration of 2 μg/mL 24 hours after transfection. Cells were harvested 24 hours post-induction for downstream analysis.

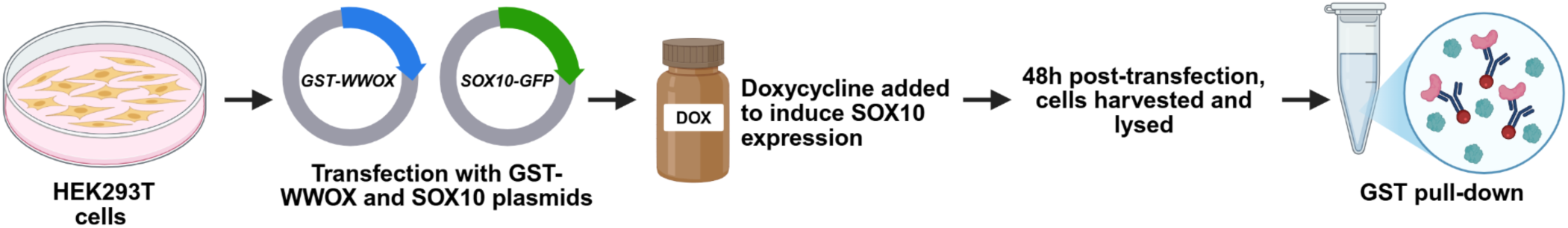

At 48 hours post-transfection, cells were collected and pelleted by centrifugation. Cell pellets were lysed in 150 μL of lysis buffer containing 0.5% NP-40, 1% phosphatase inhibitor 1 (Ph1), 1% phosphatase inhibitor 2 (Ph2), and 1% protease inhibitor cocktail (PIC). A 20 μL aliquot of the lysate was set aside as input and stored at −20 °C. The remaining 130 μL of lysate was combined with 400 μL of lysis buffer and 30 μL of GST affinity beads. The mixture was rotated on a rocker at 12 rpm for 2 hours at 4 °C. After incubation, samples were centrifuged at 4,000 rpm for 1 minute at 4 °C. The supernatant was discarded, and the beads were washed three times with 1 mL of wash buffer (lysis buffer supplemented with 1% NP-40, 1% Ph1, 1% Ph2, and 1% PIC). Beads were resuspended in 20 μL of lysis buffer and 20 μL of SDS-PAGE sample buffer. The samples were then boiled at 100 °C for 10 minutes before analysis.

#### Protein extraction from brain tissue and lysis

Frozen cortex tissue samples (25–50 mg) were lysed in 200–250 μL of lysis buffer containing 50 mM Tris (pH 7.5), 150 mM NaCl, 10% glycerol, and 0.5% Nonidet P-40 (NP-40), supplemented with protease and phosphatase inhibitors (1:100 dilution). Tissue homogenization was performed manually using a glass homogeniser to ensure complete disruption. Lysates were incubated on ice for 10 minutes, then centrifuged at 14,000 rpm for 15 minutes at 4 °C. The resulting supernatant was collected and stored at −20 °C until further analysis.

#### HA-pull down assay

For the HA pull-down assay, 25 μL (0.25 mg) of Pierce Anti-HA Magnetic Beads (Thermo Fisher, Catalog No. 88837) was transferred to a 1.5 mL microcentrifuge tube. To pre-wash the beads, 175 μL of 0.05% TBS-T was added, and the mixture was gently vortexed. The tube was placed on a magnetic stand to collect the beads, and the supernatant was discarded. The beads were washed twice with 1 mL of TBS-T, with gentle vortexing for 1 minute per wash, followed by magnetic separation and removal of the supernatant. HA-tagged protein lysates prepared from HA-Wwox mouse cortex were then added to the beads. The mixture was incubated at room temperature for 30 minutes with gentle mixing. Following incubation, the beads were collected on a magnetic stand, and the unbound fraction was removed and retained for analysis. The beads were subsequently washed three times with 300 μL of TBS-T and once with 300 μL of ultrapure water; each wash step involved gentle mixing and magnetic collection, followed by discarding of the supernatant. For protein elution, 100 μL of SDS-PAGE sample buffer was added to the beads and gently vortexed. The samples were then incubated at 95–100 °C for 5–10 minutes to denature the proteins. After incubation, the beads were separated magnetically, and the supernatant containing the eluted HA-tagged proteins was collected for downstream analysis.

#### Cycloheximide (CHX) chase assay and proteasome inhibition

To assess SOX10 protein stability, both WT and WWOX knockout Oli-neu cells were treated with 20 μg/mL cycloheximide (CHX) for 2, 4, 6, or 8 hours. For proteasome inhibition, cells were additionally treated with 10 μM MG132 in combination with CHX for 8 hours before harvesting. Following treatment, cells were lysed as previously described.

#### Ubiquitination assay on Oli-neu cells

OliNeu cells were plated in appropriate culture plates and allowed to adhere overnight. The following day, cells were transfected with 4 μg HA-Ubiquitin plasmid (Addgene plasmid #18712). Lipofectamine reagent was used at a ratio of 1μL per 1 μg of DNA, diluted in 250 μL of serum-free medium (SFM). Cells were incubated with the transfection mixture for 24 hours. 10 μM MG132 was added 4 hours before the harvesting to stabilize the proteins. For SOX10 immunoprecipitation, cell lysates were incubated with 3–5 µL of anti-SOX10 antibody at 4 °C for 1 h with gentle rotation. Subsequently, 40 µL of protein a/g plus-agarose beads (sc-2003) were added to capture the antibody–protein complexes, and the mixture was incubated overnight. at 4 °C with gentle shaking. The beads were then collected by centrifugation at 14,000 × g for 30 s at 4 °C, and the supernatant was carefully removed. The beads were washed three times with 1 mL PBS containing 25% lysis buffer and 1% PMSF, each wash performed for 10 min at 4 °C with gentle rotation. After the final wash, beads were pelleted by centrifugation (14,000 × g, 30 s, 4 °C), resuspended in 20 µL washing buffer and 5 µL 5× SDS loading buffer, and boiled at 100 °C for 5 min before Western blot analysis.

#### Immunoblotting (Western Blot)

Protein concentrations were determined, and 30–50 μg of total protein was loaded onto a 10% SDS-PAGE gel. Electrophoresis was performed at 110 V for 2.5 hours. Proteins were transferred onto a nitrocellulose membrane using a transfer system set at 25 V and 2.5 A for 15 minutes. Membranes were blocked with 5% skim milk in Tris-buffered saline with Tween-20 (TBST) for 1 hour at room temperature with gentle shaking. Following blocking, membranes were incubated with the primary antibody overnight at 4°C with gentle shaking (Rabbit Anti-WWOX 1:10000, Rabbit Anti-SOX10 (ab227680) 1:200, Rabbit anti-HSP90 (4874) 1:1000, Mouse anti-GAPDH (CB1001) 1:10000, Rabbit anti-HA (71-5500) 1:1000, Rabbit anti-GST (27457701V) 1:000). After incubation, membranes were washed three times with TBST (10 minutes each with shaking) and subsequently incubated with the appropriate HRP-conjugated secondary antibody for 1 hour at room temperature with shaking. Membranes were then washed three times with TBST (10 minutes each with shaking). Protein bands were visualised using enhanced chemiluminescence (ECL) according to the manufacturer’s instructions.

## Resource availability

### Lead contacts

Further requests for resources and reagents, data, or code should be directed to and will be fulfilled by lead contacts: Jose Davila Velderrain & Rami I. Aqeilan

### Materials availability

This study did not create new unique reagents.

### Data and Code availability

Summary information of transcriptomic data sources, processed data, and annotation tables will be available upon publication. Any additional information required to reanalyse the data is available upon request.

## Acknowledgements

The mouse oligodendrocyte progenitor cell line, Oli-neu, was a generous gift from the laboratory of Prof. Elior Peles, Department of Molecular Cell Biology, Weizmann Institute of Science. We thank the International Multiple Sclerosis Genetics Consortium for the generation of GWAS data. We thank the members of the Aqeilan’s and Davila-Velderrain’s lab for useful discussions. This study was supported by a grant from LOTTE & JOHN HECHT MEMORIAL FOUNDATION (no. 5085) to RIA. We also acknowledge the support of the Carole and Andrew Harper Diversity Scholarship Program to BAD.

## Author contributions

Conceptualisation: BAD, CM, JDV, RIA.

Methodology: BAD, CM, SAS, SR, TJ, JDV.

Investigation: BAD, CM, SAS, RA, JDV, RIA.

Visualisation: BAD, CM, SAS, JDV.

Supervision: JDV, RIA.

Writing—original draft: BAD, CM, JDV, RIA.

## Declarations of interests

R.I.A. is a consultant for Mahzi Therapeutics. The other authors declare no competing interests.

